# Constructing neural network models from brain data reveals representational transformations underlying adaptive behavior

**DOI:** 10.1101/2020.12.24.424353

**Authors:** Takuya Ito, Guangyu Robert Yang, Patryk Laurent, Douglas H. Schultz, Michael W. Cole

## Abstract

The human ability to adaptively implement a wide variety of tasks is thought to emerge from the dynamic transformation of cognitive information. We hypothesized that these transformations are implemented via conjunctive activations in *conjunction hubs* – brain regions that selectively integrate sensory, cognitive, and motor activations. We used recent advances in using functional connectivity to map the flow of activity between brain regions to construct a task-performing neural network model from fMRI data during a cognitive control task. We verified the importance of conjunction hubs in cognitive computations by simulating neural activity flow over this empirically-estimated functional connectivity model. These empirically-specified simulations produced above-chance task performance (motor responses) by integrating sensory and task rule activations in conjunction hubs. These findings reveal the role of conjunction hubs in supporting flexible cognitive computations, while demonstrating the feasibility of using empirically-estimated neural network models to gain insight into cognitive computations in the human brain.

## Introduction

The human brain exhibits remarkable cognitive flexibility. This cognitive flexibility enables humans to perform a wide variety of cognitive tasks, ranging from simple visual discrimination and motor control tasks, to highly complex context-dependent tasks. Key to this cognitive flexibility is the ability to use cognitive control, which involves goal-directed implementation of task rules to specify cognitive and motor responses to stimuli^1–3^. Previous studies have investigated how task-relevant sensory, motor, and rule features are represented in the brain, finding that sensory stimulus features are represented in sensory cortices^4, 5^, motor action features are represented in motor cortices^6^, while task rule features are represented in prefrontal and other association cortices^3, 7–10^. However, these studies focused on *where* cognitive representations are located in the brain, rather than *how* the brain uses and transforms those representations^11^. For example, during context-dependent tasks, exactly how the brain converts incoming sensory stimulus activity into motor activity remains unclear^12^. In contrast, artificial neural network models (ANNs) can provide computationally rigorous accounts of how context and stimuli input vectors interact to perform complex tasks^13, 14^. Inspired by the formalization of ANNs, we show how task rule and sensory stimulus activations are transformed into motor response activations in the human brain via intrinsic functional connectivity (FC) weights. We achieve this by constructing an empirically-estimated neural network (ENN) model from fMRI data to provide insight into the neural transformations in the brain during a cognitive control task.

The Flexible Hub theory provides a network account of how large-scale cognitive control networks implement flexible cognition by updating task rule representations^15, 16^. While the Flexible Hub theory primarily focuses on the importance of flexible rule updating for complex task performance, it does not specify how rules interact with incoming sensory stimulus activity. However, the Flexible Hub theory was built upon the Guided Activation Theory of prefrontal cortex – a seminal theory of the neural correlates underlying cognitive control – which posits that successful performance of a cognitive control task requires the selective mixing of task context with sensory stimulus activity^3^. The selective mixing of task context and sensory stimulus activations would produce conjunctive activations that implement task rules on sensory stimuli. Conjunctive activations refer to task-related activations that represent the conjunction (binding) of multiple different task conditions, such as task rules and sensory stimuli^9, 12^. For example, an activation representing the conjunction of rule X and stimulus Y could be active only when stimulus Y is presented with rule X, not when stimulus Y is presented without rule X. These conjunctive activations are thought to form through inter-area guided activations in brain areas hidden somewhere in association cortex, which we term *conjunction hubs* (Fig. 1a). The outputs of conjunction hubs then generate motor activations to produce task-appropriate behavior. Thus, by testing the hypothesis put forth in the Guided Activation Theory of interacting rule- and stimulus-guided neural activations (i.e., conjunctions)^3, 17^, we built upon the Flexible Hub theory to provide insight into flexible task control.

**Figure 1.**
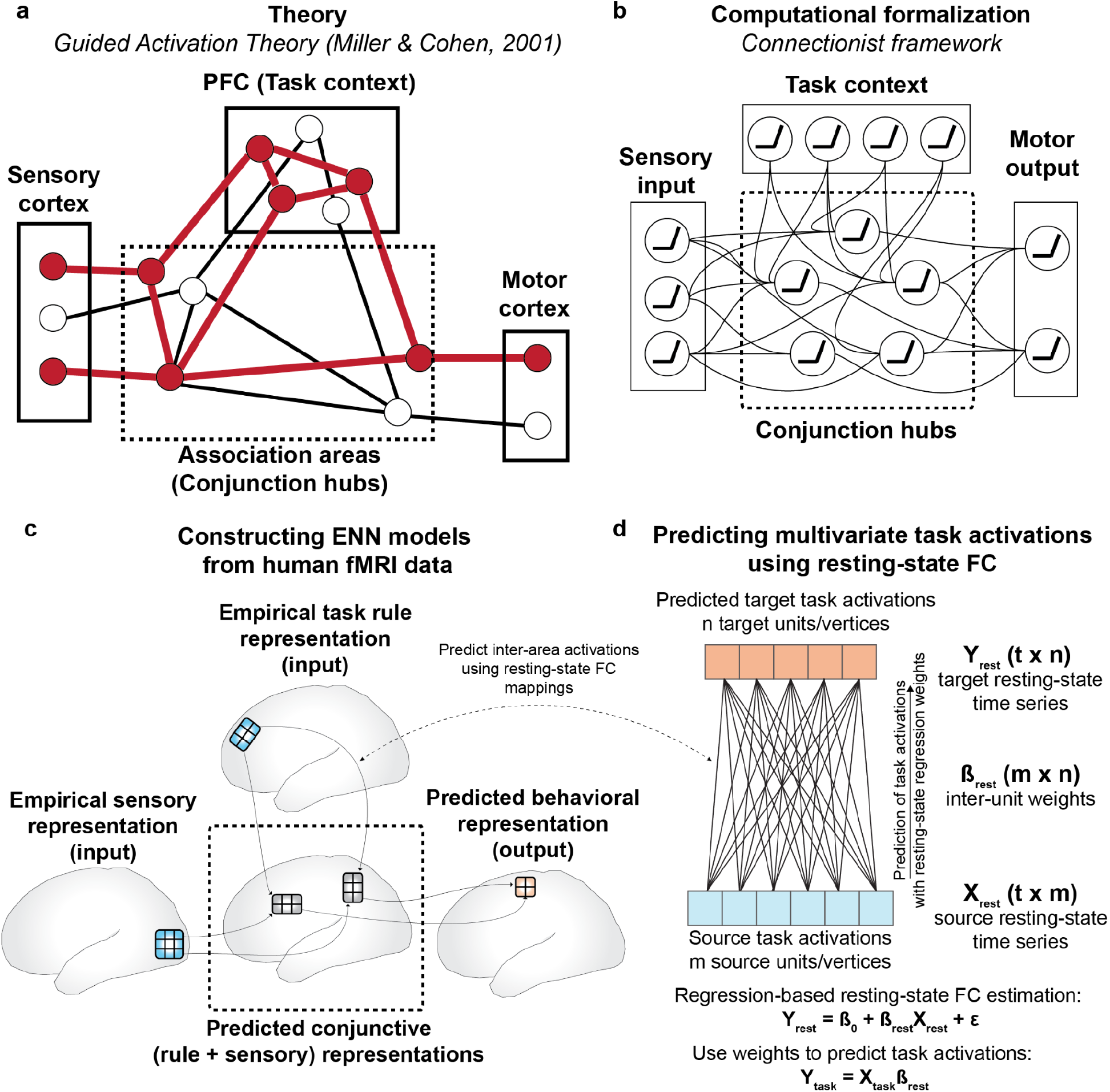
Leveraging the Guided Activation Theory to inspire ENN models of cognitive computation during task-based fMRI. **a)** A modified version of the Guided Activation Theory of prefrontal cortex, highlighting a potential key role for conjunction hubs. The Guided Activation Theory posits that sensory cortices (left), which contain sensory stimulus-related activations, and prefrontal areas (top), which contain task context/rule activations, integrate in association cortex to produce conjunctive activations through patterns of guided activations. Conjunctive activations are then guided to motor areas to generate motor response activations for task behavior. **b)** The Guided Activation Theory can be reconceptualized in a connectionist framework. This provides a formalization of how flexible sensorimotor transformations may be implemented computationally. The formalization involves the task context and sensory stimuli representing the input layer, the association units representing a hidden layer, and the behavioral (motor) responses as the output layer. **c)** Testing the Guided Activation Theory using task fMRI data collected in humans during context-dependent tasks. Using quantitative methods, we empirically test how different task activations (e.g., sensory stimuli and task context) form conjunctive activations to produce motor response activations using activity flow mapping^18^. **d)** The Guided Activation Theory can be empirically tested by projecting task activation patterns between brain areas by estimating inter-area FC weight mappings obtained from resting-state fMRI data. Based on the activity flow principle^18^, we estimated inter-vertex mappings using regression (see Methods) on resting-state fMRI data. This approach identifies a projection that maps across distinct spatial units (i.e., vertices) in empirical data, similar to how inter-layer weights propagate activity across layers in an ANN.

We recently developed a method – activity flow mapping – that provides a framework for testing the Guided Activation Theory with empirical brain data^18^. Activity flow mapping involves three steps. First, a network model is derived from empirically-estimated connectivity weights. Second, empirical task activations (e.g., activity patterns from sensory regions) are used as inputs to simulate the activity flow (i.e., propagating activity) within the brain network model. Finally, the predictions generated by simulated activity flow are tested against independent empirical brain activations for model validation. Here we used activity flow mapping to test whether empirical task activations and FC could model transformations from sensory stimulus activations to motor response activations during a context-dependent cognitive paradigm.

We sought a principled approach to identify brain areas that form the conjunctive activations hypothesized to produce flexible behavior. Recent studies have successfully used trained ANNs to identify cognitive representations during tasks^13, 14, 19^. Importantly, the representations of ANNs have often converged with representations found in neural data^20–22^, suggestive of the utility of ANNs in investigating task representations in the brain. Inspired by these previous studies, we first constructed a simple ANN to investigate how conjunctive representations formed from task context and stimulus input activations during a 64-context cognitive paradigm. Using a simple ANN trained to perform the same task allowed us to identify putative conjunctive representations within the ANN that integrated rule and stimulus activations. This provided a blueprint to search for similar representations in our human brain data. After identifying the representation of task context and stimulus conjunctions in the ANN, we identified brain regions – conjunction hubs – with similar conjunctive representations in fMRI data. The identification of brain regions selective for task rules, sensory stimuli, motor responses, and conjunctions, made it possible to construct an ENN (which is derived from brain data and distinct from the ANN) and empirically test the Guided Activation Theory with activity flow mapping over data-derived functional connections. We found that behavioral activations (in motor cortices) could be predicted through the formation of conjunctive activations through activity flow guided by task rule and sensory stimulus activations.

To summarize, we provide an empirical demonstration of connectionist-style computations in fMRI data during a 64-context cognitive paradigm. This was achieved by constructing a task-performing ENN directly from fMRI data, empirically testing the plausibility of connectionist-like computations conceptualized by the Guided Activation Theory. Importantly, the original conceptualization of the Guided Activation Theory did not specify an exact implementation in neural data. Thus, in this study we identify specific components of an ENN – a brain-based connectionist model (e.g., brain regions and connectivity weights) – that were critical for implementing context-dependent representational transformations, while also revealing corresponding failure modes (e.g., alternative connectionist models that failed to transform representations). This involved identifying the brain areas selective to different task components, namely task rules, sensory stimuli, motor responses, and conjunctions. These areas formed the spatial areas/layers of the ENN, which are conceptually similar to layers in an ANN. Next, in contrast to ANNs, which typically use supervised learning to estimate connectivity weights between layers, we show that activations in ENNs can be transformed via activity flow over FC weights estimated from resting-state fMRI (Fig. 1d). This resulted in a task-performing ENN model that transforms stimulus and task-rule fMRI activations into response activations in motor cortex during a flexible cognitive control task. Critically, the transformations implemented by the ENN were carried out without classic optimization approaches such as gradient learning, demonstrating that the intrinsic architecture of the resting brain is suitable for implementing representational transformations. Together, these findings illustrate the computational relevance of functional network organization and the importance of conjunctive representations in supporting flexible cognitive computations in the human brain.

## Results

### Identifying brain areas containing task-relevant activations

The Flexible Hub Theory posits that rapid updates to rule representations facilitate flexible behavior^15, 16^, while the Guided Activation Theory^3^ states that sensory stimulus and task rule activations integrate in association cortex to form conjunctive activations (Fig. 1a,c). Thus, due to its comprehensive assessment of rule-guided sensorimotor behavior across 64 task contexts, we used the Concrete Permuted Rule Operations (C-PRO) task paradigm^5^ to test both theories (Fig. 2a). Briefly, the C-PRO paradigm is a highly context-dependent cognitive control task, with 12 distinct rules that span three rule domains (four rules per domain; logical gating, sensory gating, motor selection). These rules were permuted within rule domains to generate 64 unique task contexts, and up to 16384 unique trial possibilities (with various stimulus pairings; see Methods). We chose this cognitive paradigm largely due to its systematic use of counterbalancing of task elements (stimuli, contexts, and responses) across trials, which allowed us to rigorously separate the motor response activations from the sensory and context cue activations (due to careful counterbalancing and averaging; Supplementary Fig. 9).

**Figure 2.**
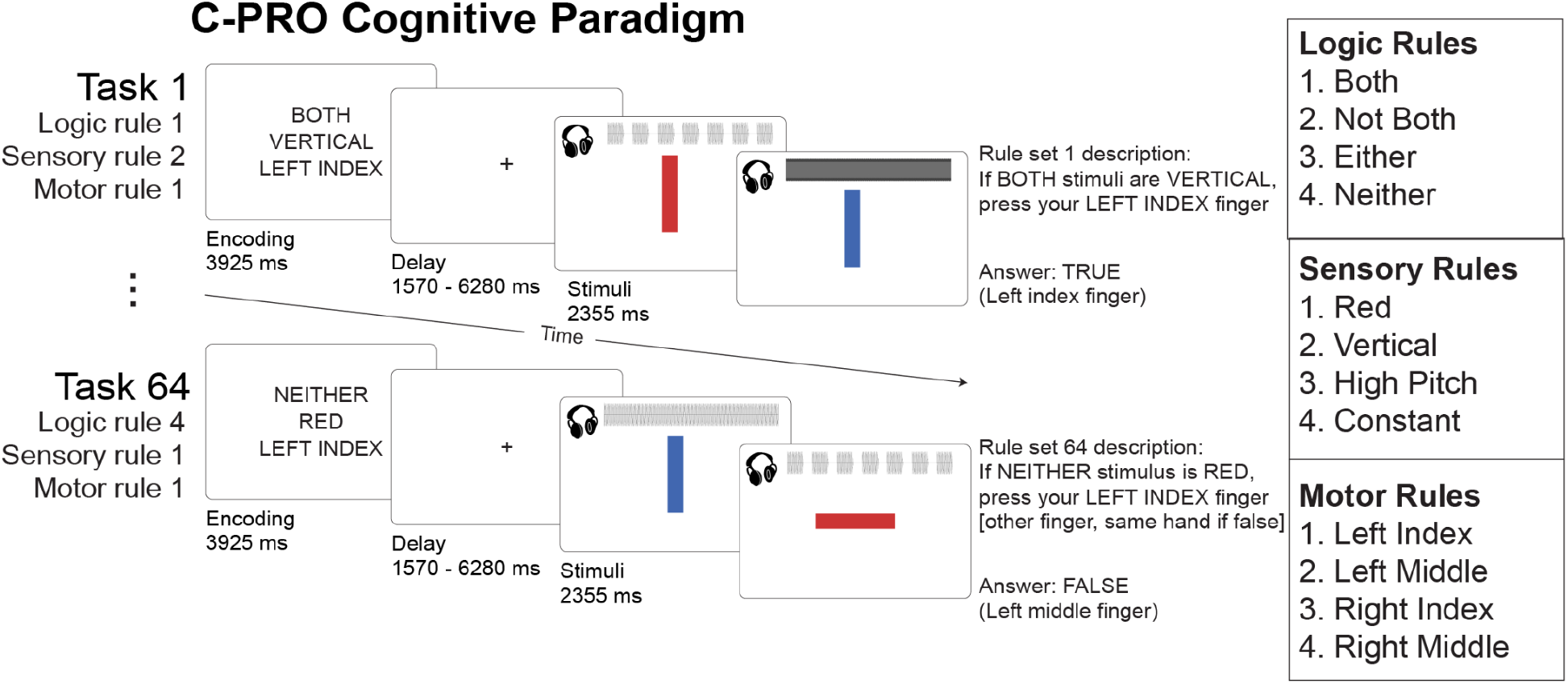
The Concrete Permuted Rule Operations (C-PRO) task paradigm^8^. For a given trial, subjects were presented with a task rule set (context), in which they were presented with three rules sampled from three different rule domains (i.e., logical gating, sensory gating, and motor selection domains). After a delay period, subjects applied the task rule set to two consecutively presented sensory stimuli (simultaneous audio-visual stimuli) and responded accordingly with button presses (index and middle fingers on either hand). We employed a miniblock design, in which for a given task rule set, three stimulus periods were presented separated by an inter-trial interval (1570ms). See Methods for additional details.

To test both the Flexible Hub and the Guided Activation theories, we needed to identify the set of regions responsive to different task components (sensory stimuli, task context, motor responses, and conjunctions). We first identified the set of cortical areas that contained decodable sensory stimulus activations (Fig. 3a). Because our stimuli were multimodal (audiovisual), this involved the identification of surface vertices that contained the relevant visual (color and orientation) and auditory (pitch and continuity) dimensions. We performed a four-way classification^23^ (using a minimum-distance/nearest-neighbor classifier^24^) to decode stimulus pairs for each of the four stimulus dimensions (e.g., red-red vs. red-blue vs. blue-red vs. blue-blue). Decoding analyses were performed within each brain parcel using the Glasser et al. atlas^25^, using vertices within each parcel as decoding features. For all decoding analyses, statistical thresholding was performed using a one-sided binomial test (greater than chance=25%), and corrected for multiple comparisons using an FDR-corrected p<0.05 threshold. We collectively defined the units in the ENN (i.e., vertices) that contained sensory stimulus activity to be the set of all vertices within the parcels that contained decodable stimulus activity (Fig. 3b; Supplementary Tables 1-4).

**Figure 3.**
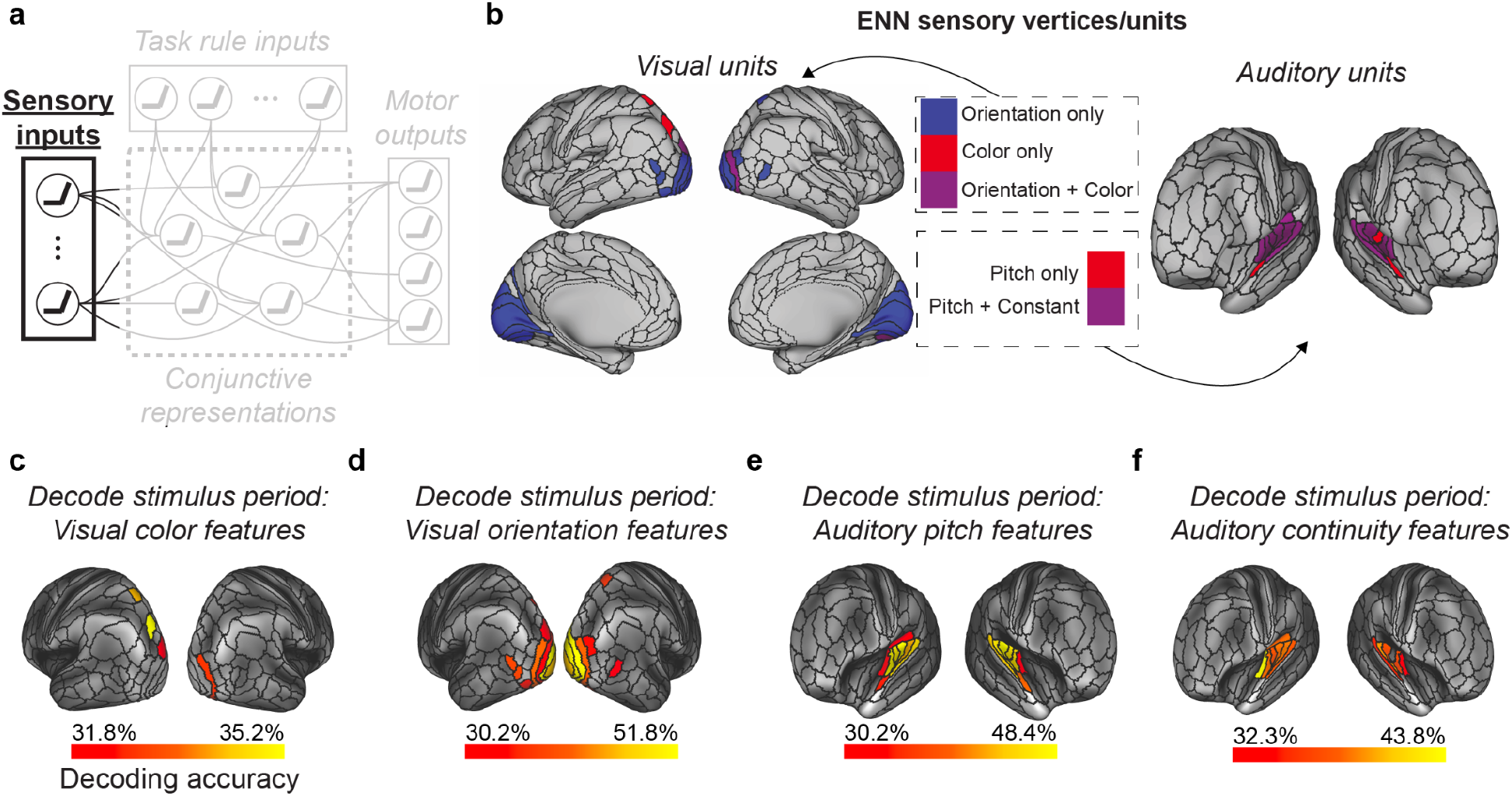
Identifying sensory stimulus input units (vertices) of the ENN using an fMRI decoding analysis. **a)** We identified the sensory stimulus representations in empirical data using fMRI pattern decoding of stimulus activations. This corresponded to the sensory input component of the Guided Activation Theory. To decode visual features (i.e., color and orientation stimulus features) we decoded the vertices within each parcel in the visual network using a recent functional network atlas^26^. To decode auditory features (i.e., pitch and continuity) we decoded the vertices within each parcel in the auditory network (see Methods). **b)** The ENN sensory units, which were derived from a mask of the vertices that could successfully decode stimulus features (panels **c-f**). **c)** Decoding of color features using task activation estimates (from a task GLM) during the stimulus presentation period of the C-PRO task. Chance was 25%; cortical maps were thresholded using an FDR-corrected threshold of p<0.05. **d)** 4-way decoding of orientation features. **e)** 4-way decoding of auditory pitch features. **f)** 4-way decoding of auditory continuity features.

Next, we performed a 12-way decoding analysis – isolated to the fMRI activation during the task encoding period – across all 12 task rules to identify the set of vertices that contained task rule activity. Our previous study illustrated that rule representations are widely distributed across cortex^8^, such that we tested for rule representations in every parcel in the Glasser et al. atlas (360 total parcels^25^). In addition, another study in non-human primates found that sensory areas also contain high-level task rule information, likely due to top-down feedback from higher-order areas^27^. Consistent with these findings, we again found that task rule representations were widely distributed across cortex (Fig. 4b; FDR-corrected p<0.05 threshold; Supplementary Table 6). The set of vertices that survived statistical thresholding were included as “task rule” input units in the ENN (Fig. 1c). Since rule representations were widely distributed across cortex, we next quantified the contribution of each rule activation in predicting conjunctive activations. We found that while both sensorimotor and association networks had similarly high levels of activations during the task rule encoding period (Supplementary Fig. 2a,b), many of these activations were dampened by their FC to conjunctive areas. Activations that contributed most to conjunctive activations were primarily from the dorsal attention network (Supplementary Fig. 2c,d). These results suggest that despite widespread task rule activations, some regions (e.g., dorsal attention network) played a disproportionate role in task rule implementation and the selection of conjunctions.

**Figure 4.**
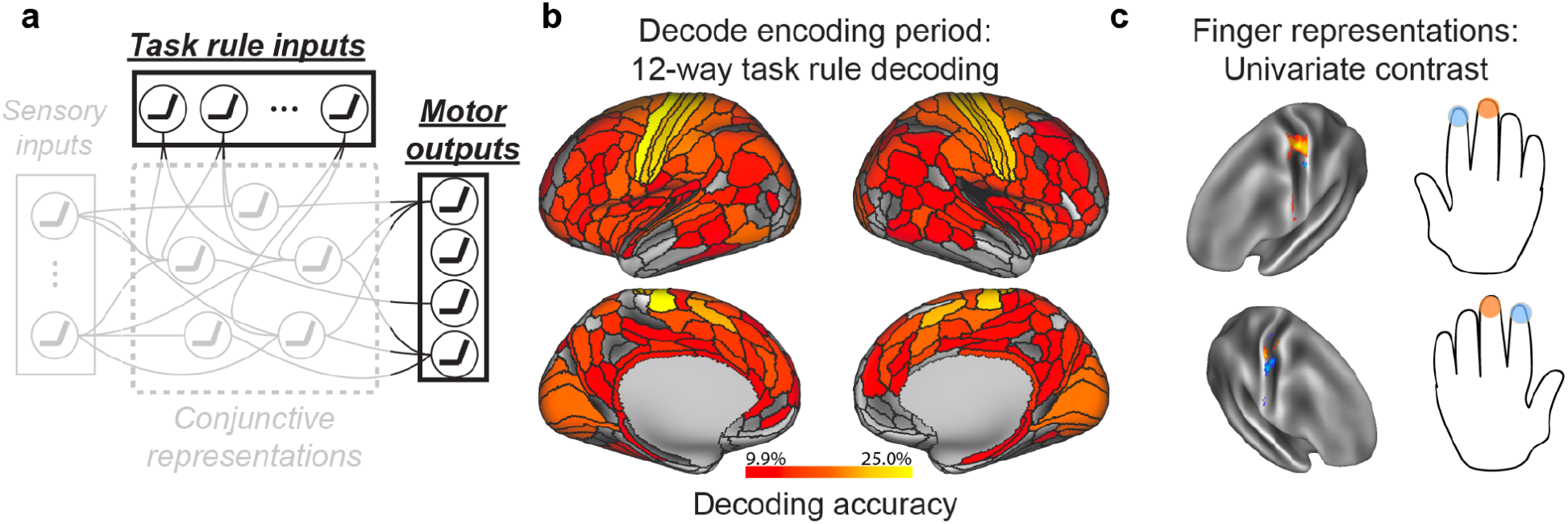
Identifying ENN units (i.e., fMRI vertices) containing relevant task rule (context) and motor response (behavior) representations. **a)** We identified the task rule input and motor output representations in empirical data using MVPA and univariate task activation contrasts. **b)** A 12-way decoding of each of the task rules (across the 3 rule domains) using task activations (estimated from a task GLM) during the encoding period of the C-PRO task. We applied this 12-way decoding to every parcel, given that task rule activations have been previously shown to be widely distributed across cortex^8^. Chance decoding was 8.33%; statistical maps were thresholded using an FDR-corrected p<0.05 threshold. **c)** To identify the motor/output activations, we performed a univariate contrast, contrasting the middle versus index finger response activations for each hand separately. Finger response activations were estimated during the response period, and univariate contrasts were performed on a vertex-wise basis using all vertices within the somatomotor network^26^. Contrast maps were statistically thresholded using an FDR-corrected p<0.05 threshold. The resulting finger activations matched the placement of finger representations in the well-established somatomotor homunculus in the human brain.

The C-PRO task paradigm required button presses (using index and middle fingers on either hand) to indicate task responses. We were able to take advantage of well-established knowledge of the localization of these finger representations in primary motor cortex^28^, rather than conducting a large search for representation of the relevant information (e.g., as we did for task rules). This had the advantage of putting the ENN to a more stringent test; requiring the ENN to select representations of motor responses in the format known to directly cause the processes of interest (i.e., increased neural activity in M1 finger representations causing motor behavior). Thus, to isolate finger representations in empirical fMRI data, we performed a univariate contrast of the vertex-wise response-evoked activation estimates during index and middle finger response windows (see Methods). We performed univariate analyses rather than multivariate decoding analyses for motor response identification since there were only two conditions – index and middle finger responses – to distinguish and because (unlike the other functional localizations) we knew the direction of amplitude change (increased activity) throughout the localized motor representations. For each hand, we performed a two-sided paired t-test (paired across subjects) for middle versus index finger responses in M1/S1 parcels. Contrast maps were corrected for multiple comparisons (comparisons across vertices) using an FDR-corrected threshold of p<0.05 (Fig. 4c). Vertices that survived statistical thresholding were then selected for use as output units in the ENN (Fig. 1c).

### Identifying conjunction hubs

We next sought to identify conjunctive representations that could plausibly implement the transformation of input to output activations across the 64 task contexts (Fig. 5a). However, we were uncertain as to what sorts of activation patterns (i.e., representations) we would expect in putative conjunction hubs. Thus, we began by building an ANN that formalizes the Guided Activation Theory (Fig. 1b). We trained the ANN model on an analogous version of the C-PRO task until the model achieved 99.5% accuracy (see Methods). We were specifically interested in characterizing the representations in the hidden layers, since these activations necessarily integrated task rule and sensory stimulus activations (i.e., conjunctions). To identify the task rule and sensory stimulus conjunctive representations, we performed a representational similarity analysis (RSA) on the hidden layers of the ANN^24^. The representational similarity matrix (RSM) of the hidden layers consisted of 28 task activation features: 12 task rules (which spanned the 3 rule domains), and 16 stimulus pairings (which spanned each sensory dimension). We then compared the RSM of the ANN’s hidden units (Fig. 5b) to RSMs of each brain region in the empirical fMRI data (Fig. 5c). This provided a map of brain regions with similar representations to those of the ANN’s hidden units, which contain the conjunction of task rule and sensory stimulus activations.

**Figure 5.**
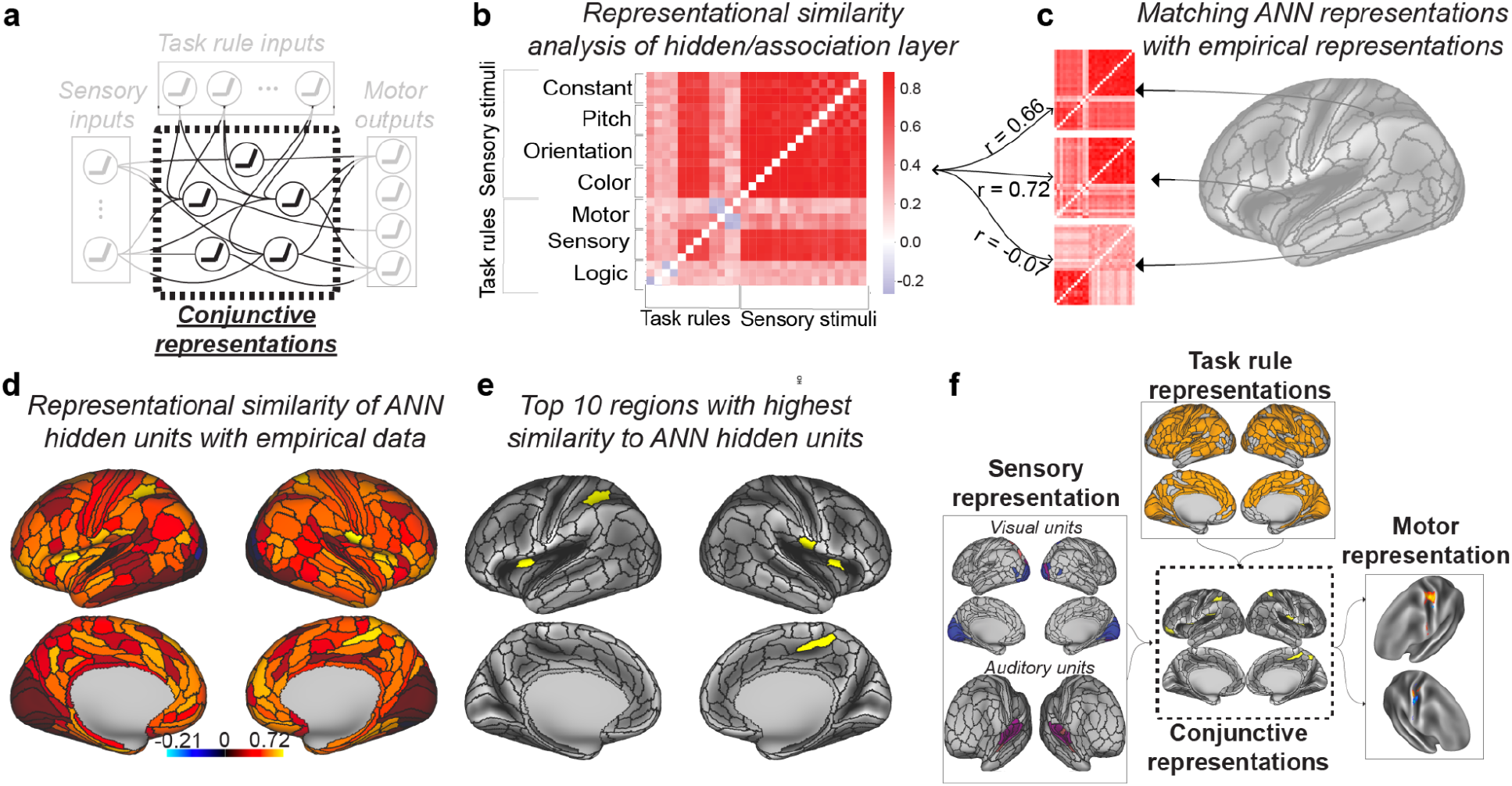
Identifying conjunction hubs: brain areas (vertices) that contain task-relevant conjunctions of sensory stimulus and task rule activations. **a)** The Guided Activation Theory states that there exist a specific set of association (or hidden) areas that integrate sensory stimulus and task context activations to select appropriate motor response activations. In an ANN where task rules and sensory stimulus activations serve as inputs, the ANN’s hidden layers integrate rule and stimulus activations providing a computational framework that is analogous to the role suggested for association areas in the Guided Activation Theory. **b)** We therefore used the representational similarity matrix (RSM) of the ANN’s hidden layers as a blueprint to identify analogous conjunctive activations in empirical data. **c)** We constructed RSMs for each brain parcel (using the vertices within each parcel as features). We evaluated the correspondence between the representational geometry of the ANN’s hidden layers and each brain parcel’s representational geometry. Correspondence was assessed by taking the correlation of the upper triangle of the ANN and empirical RSMs. **d)** The representational similarity of ANN hidden units and each brain parcel. **e)** We showed the top 10 regions with highest similarity to the ANN hidden units. **f)** The full ENN architecture for the C-PRO task. We identified the vertices that contained task-relevant rule, sensory stimulus, conjunctive, and motor output activations.

To evaluate the similarity of the ANN’s hidden representational geometry with each brain parcel, we computed the similarity (using Spearman’s correlation) of the ANN’s RSM with the brain parcel’s RSM (Fig. 5c). This resulted in a cortical map, which showed the similarity between each brain region and the ANN’s hidden representations (Fig. 5d). For our primary analysis, we selected the top 10 parcels with highest similarity to the ANN’s hidden units to represent the set of spatial units that contain putative conjunctive activations in the ENN (Fig. 5e). The conjunction hubs were strongly represented by the cingulo-opercular network, a network previously reported to be involved in task set maintenance and a variety of other cognitive control functions (Supplementary Fig. 3; Supplementary Table 5)^29^. However, other association networks also had strong associations with the ANN’s hidden layer representations (Supplementary Fig. 3b). We also performed ENN simulations using the top 20, 30, and 40 regions with highest similarity to the ANN hidden units (see text below). To ensure that the RSM identified from the ANN was critical to perform the task, we further identified a control ANN’s RSM, where we shuffled all parameters within each layer after training the model (see Methods). We found that in addition to the model no longer performing the task correctly, the model contained little representational structure (no representational dissimilarities across conditions) (Supplementary Fig. 4). The control ANN’s hidden layer also had significantly weaker similarity to the empirical RSMs at each parcel (Supplementary Fig. 4d).

### Task-performing neural network simulations via empirical connectivity

The previous sections provided the groundwork for constructing an ENN model from empirical data. After estimating the FC weights between the surface vertices between ENN layers using resting-state fMRI (see Methods), we next sought to evaluate whether we could use this ENN to produce representational transformations sufficient for performing the C-PRO paradigm. This would demonstrate that the empirical input activations (task rule and sensory stimulus activations) and the estimated connectivity patterns between ENN layers are sufficient to approximate the cognitive computations involved in task performance.

The primary goal was to generate a motor response activation pattern (i.e., behavior) that we could then compare to correct task performance. The only inputs to the model were a combination of activation patterns for a specific task context (rule combination) and sensory stimulus pair sampled from empirical data (Fig. 6a), which we term “pseudo-trials”. (“Pseudo-trials” refer to simulated trials using estimated activations rather than the actual experimental trials subjects performed.) The outputs of the model were the predicted motor response activation pattern in motor cortex that should correspond to the correct button press (Fig. 6c). High correspondence between the predicted and actual motor activation patterns would constitute an empirical identification of representational transformation in the brain, where task rule and sensory stimulus activity is transformed into task-appropriate response activation patterns in motor cortex.

**Figure 6.**
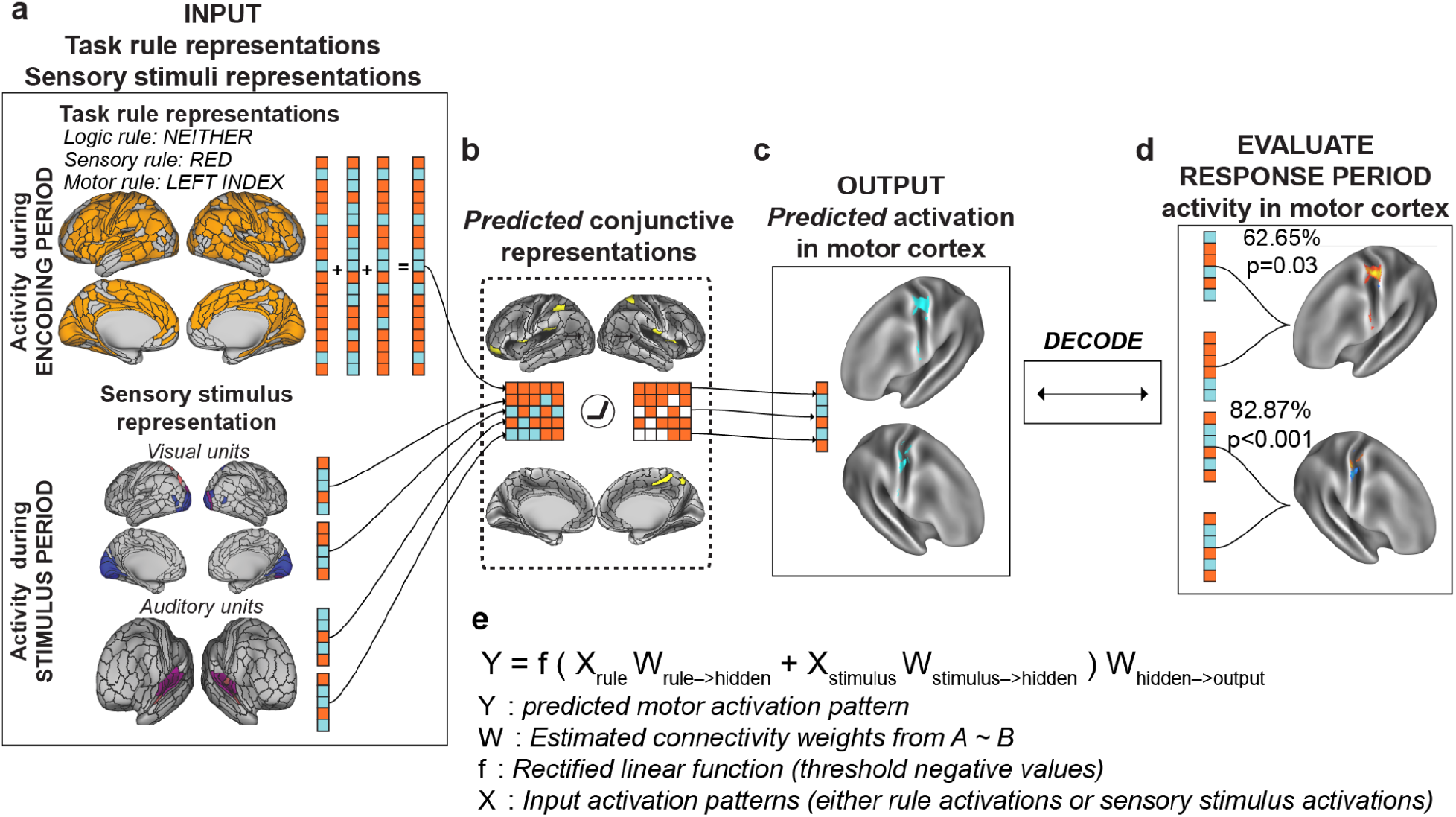
Simulating context-dependent sensorimotor transformations with empirically-estimated task activations and inter-unit FC estimates. We constructed the ENN by identifying the vertices that contained task rule, sensory stimulus, and motor response activations and by estimating the resting-state FC weights between them. **a)** The input layer, consisting of vertices with decodable task rule and sensory stimulus activations. **b)** Through activity flow mapping, input activations were mapped onto surface vertices in conjunction hubs. The activity flow-mapped vertices were passed through a nonlinearity, which removed any negative values. This threshold was chosen given the difficulty in interpreting *predicted* negative BOLD values. **c)** The predicted conjunctive activations were then activity flow-mapped onto the motor output vertices, generating a predicted motor activation pattern. **d)** These predicted motor activations were then tested against the actual motor response activations of other subjects using a leave-8-subject out cross validation scheme. A decoder was trained on the predicted motor response activations and tested on the actual motor response activations of the held-out cohort (see Methods and Supplementary Fig. 1). **e)** An equation summarizing the ENN model’s computations.

Simulating activity flow in the ENN involved first extracting the task rule activation patterns (inputs) for a randomly generated task context (see Methods and Supplementary Fig. 1). Independently, we sampled sensory stimulus activation patterns for each stimulus dimension (color, orientation, pitch, continuity) (Fig. 3). Then, using activity flow mapping with resting-state FC weights, we projected the activation patterns from the input vertices onto the conjunction hub vertices (Fig. 6b). Supplementary Fig. 2 provides a visualization of the contributing vertices (via activity flow mapping) from the task rule layer onto the conjunction hubs, finding that despite widespread task rule activations across most of cortex, the dorsal attention network plays a disproportionate role in generating conjunctive activations (Supplementary Fig. 2d). The predicted conjunction hub activation pattern was then passed through a simple rectified linear function, which removed any negative values (i.e., any values lower than resting-state baseline; see Methods). Thresholded values were then projected onto the output layer vertices in motor cortex (Fig. 6c), yielding a predicted response activation pattern. The sequence of computations performed to generate a predicted motor activation pattern (Fig. 6a-c) is encapsulated by the equation in Fig. 6e. Thus, predicted motor activation patterns can be generated by randomly sampling different task context and sensory stimuli activations for each subject.

While the above procedure yielded a predicted activation pattern in the motor output layer, these predictions may not actually yield meaningful activation patterns. Thus, we evaluated whether the model-generated motor activation patterns accurately predicted the *actual* motor response activation patterns extracted (via GLM) during subjects’ response period. Activity flow simulations using only input task activations from the task encoding period and stimulus presentation period (Fig. 6a) generated predicted motor responses for each subject (Supplementary Fig. 1). Using a leave-8-subjects out cross-validation scheme, we trained a decoder on the four possible predicted motor responses and decoded the four possible actual motor responses (Fig. 6c,d). Training a decoder on the predicted activations and decoding the actual activations (rather than vice versa) made this analysis more in line with a prediction perspective – we could test if, in the absence of any motor task activation, the ENN could predict actual motor response activation patterns that correspond to correct behavior. We averaged motor response patterns across pseudo-trials to yield four predicted motor response activations per subject. This averaging eliminated the possibility of any remaining rule and sensory information being present in the model-generated motor response patterns given that pseudo-trials were perfectly counterbalanced across contexts, stimuli, and response.

We note that this decoding analysis is highly non-trivial, given that the predicted motor responses (which are generated from task rule and stimulus activations) are tested against the true motor responses of held-out subjects. By simulating neural network computations from stimulus and task context activations to predict motor response, we accurately decoded the correct finger response on each hand separately: decoding accuracy of right hand responses = 62.65%, non-parametric p=0.03; decoding accuracy of left hand responses = 77.58%, non-parametric p<0.001. These results demonstrate that task rule and sensory stimulus activations can be transformed into motor output activations by simulating multi-step neural network computations using activity flow mapping on empirical fMRI data. In the following sections, we illustrate that multiple control and lesion models severely impair model performance, suggesting that the constructed model contained 1) no biases towards predicting motor responses, and 2) provided the sufficient features to implement context-dependent sensorimotor transformations.

In addition, a good test of the non-triviality of the predicted motor responses would be to ensure that motor responses cannot be linearly decoded from the input activations (task rule and stimulus activations). Here, we establish that linear decoding of motor responses using input activations fails under the current decoding scheme, given that predicted activations are averaged across pseudo-trials for each motor response (see Methods). While specific task context and stimulus combinations produce a motor response at the trial level, averaging across completely counterbalanced inputs for each response leads to activations that are mathematically identical. For example, both the left index and left middle finger motor responses were averaged across red-red, red-blue, blue-red, and blue-blue stimulus events, yielding identical activity patterns across those two motor responses for all regions representing visual rather than motor information. (The same logic can be applied for all task rules.) This makes it impossible for a linear decoder to learn mappings between inputs and responses when averaging inputs, but possible for a linear decoder to learn mappings on the outputs. This is because the outputs were generated via a nonlinear function (i.e., the ENN) applied at the trial level and then averaged across trials for the decoder. We verified this empirically, finding that the accuracy was at chance, since the decoder could not classify identical inputs.

We observed an overall difference in the ability to decode left versus right hand sensorimotor transformations. However, this discrepancy was also observed when decoding actual motor response activations (rather than predicted activations) during the response period, suggesting this was an intrinsic property of the fMRI data we used (rather than the ENN) and/or due to differences in the number of identified vertices associated with response on either hand (Fig. 7h,i).

**Figure 7.**
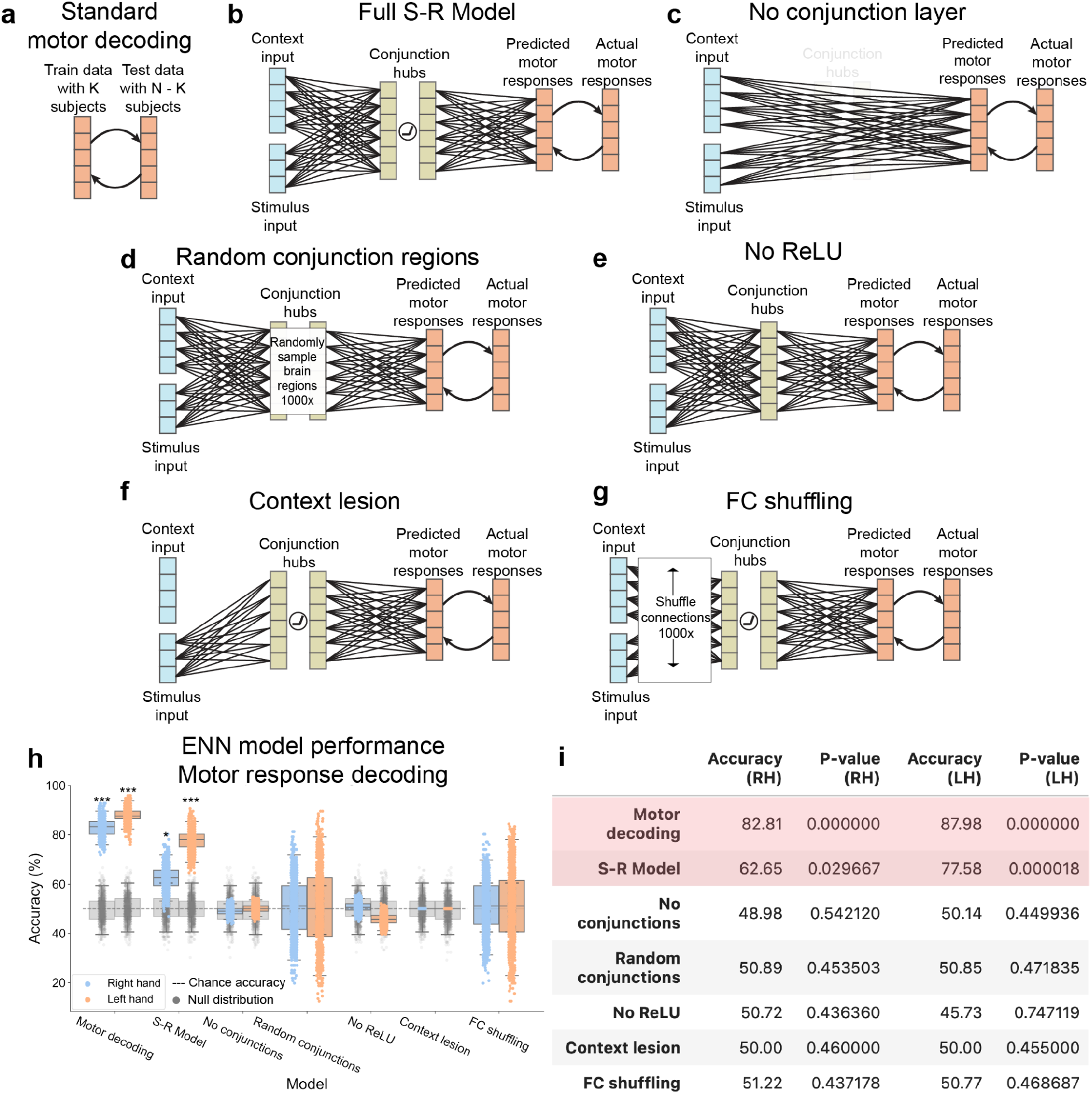
Systematic alteration of ENN model architecture verifies validity of “full S-R model” results. **a)** We first benchmarked the motor response decoding accuracy for each hand separately using a standard cross-validation scheme on motor activation patterns for each hand (tested across subjects). This standard motor decoding was done independently of modeling sensorimotor transformations. **b)** The full stimulus-response model, taking stimulus and context input activations to predicting motor response patterns in motor cortex. **c)** The ENN model after entirely removing the conjunction hubs. **d)** The ENN model, where we randomly sampled regions in the hidden layer (conjunction hubs) 1000 times and estimated task performance. **e)** The ENN model after removing the nonlinearity (ReLU) function in the hidden layer. **f)** The ENN model after lesioning connections from the task context input activations. **g)** The ENN model, where we shuffled the connectivity patterns from the stimulus and context layers 1000 times. **h)** Benchmarking the performances of all model architectures. Accuracy distributions were obtained by running 1000 iterations of the same cross-validation scheme (leave-4-out cross-validation scheme and randomly sampling within the training set; see Methods for clarification). For each iteration, we calculated a p-value, and then averaged all p-values. Boxplot whiskers reflect the 95% confidence interval. Grey distributions indicate the null distribution generated from permutation tests (permuting labels 1000 times). (*** = p<0.001; ** = p<0.01; * = p<0.01) **i)** Summary statistics of model performances. Reported accuracy is the mean across the distribution.

### The importance of the conjunctive representations

We next evaluated whether specific components of the ENN model were necessary to produce accurate stimulus-response transformations. We first sought to evaluate the role of the conjunction hubs (hidden layer) in ENN performance. This involved re-running the ENN with the conjunction hubs removed (Fig. 7c), which required resting-state FC weights to be re-estimated between the input and motor output regions directly. We found that the removal of conjunction hubs severely impaired task performance to chance accuracy (RH accuracy=48.98%, p=0.54; LH accuracy=50.14%, p=0.45; Fig. 7h,i). This illustrated the importance of conjunction hub computations in producing the conjunctive activations required to perform context-dependent stimulus-response mappings^15^.

We next replaced conjunction hubs with randomly sampled parcels in empirical data. This assessed the importance of using the ANN’s hidden layer RSM to identify conjunction hubs in fMRI data (Fig. 7d). We sampled random parcels 1000 times, recomputing the vertex-wise FC each time. The distribution of randomly selected conjunction hubs did not yield task performance accuracies that were statistically different than chance for both hands (RH mean accuracy=50.89%, p= 0.45; LH mean accuracy 50.85%, p=0.47; Fig. 7h,i). However, the overall distribution had high variance, indicating that there may be other sets of conjunction hubs that would yield above-chance (if not better) task performance. However, compared to the conjunction hubs we identified by matching empirical brain representations with ANN representations, we found that the ANN-matched conjunction hubs performed better than 83.3% of all randomly selected conjunction hubs for RH responses, and greater than 96.4% of all randomly selected conjunction hubs for LH responses.

In addition, we evaluated whether the precise number of hidden regions was critical to task performance. We ran the full ENN model, but instead of using only the top 10 regions with highest similarity to the ANN’s hidden layer’s representations, we constructed ENN variants containing the top 20, 30, and 40 hidden regions. We found that we were able to reproduce correct task performance using 20 hidden regions (RH accuracy=63.90%, p<0.001; LH accuracy=76.95%, p<0.001). Using 30 hidden regions yielded reduced yet above-chance accuracies for RH responses, but not for LH responses (RH accuracy=59.83%, p=0.024; LH accuracy=43.54%, p=0.917). Inclusion of an additional 10 hidden regions (totaling 40 hidden regions) did not yield above-chance predictions of motor responses for either hand. Inclusion of additional intermediate conjunctive regions likely introduced additional noisy (or irrelevant) activations that in turn degraded the final predicted motor output activations from which we decoded. These results suggest that conjunction hubs were better identified the greater the similarity of a region’s representational geometry was to that of the ANN’s hidden layer.

### The importance of nonlinearities when combining rule and stimulus activations

We next tested the impact of removing the rectified linear (ReLU) nonlinear functions in the conjunction hubs. This is equivalent to removing nonlinearities in an ANN, which is well-known to eliminate the ability of ANNs to implement conditional logic (e.g., the XOR logic gate)^17, 30, 31^. Conditional logic was essential to all of the C-PRO task sets, since the motor response must be chosen conditional on both the stimulus information and task rule information. Consistent with the expected computational role of the ReLU nonlinearity, we found that the removal of the ReLU function substantially impaired model performance (RH accuracy=50.72%, p=0.44; LH=45.73%, p=0.75; Fig. 7h and 7i). This is due to the fact that context-dependent sensorimotor transformations require nonlinear mappings between stimuli and responses, as predicted by the biased competition theory and validated by prior computational studies^17, 31, 32^. To more rigorously assess the impact of the ReLU on the activity flow-predicted conjunctive region representations, we compared the conjunctive representations of the full ENN model (ReLU included), ReLU removed ENN model, and the actual fMRI activations of the conjunction regions to the representations found in the ANN’s hidden layer. (Comparison of representations was measured as the cosine similarity of the representational similarity matrices.) As expected, we found that compared to the activity flow-predicted representations without the ReLU (cosine=0.44), the full ENN with the ReLU had higher similarity to the ANN’s hidden layer (cosine=0.60) (Supplementary Fig. 5). This suggested that the ReLU supports accurate context-dependent sensorimotor transformations by producing internal conjunctive representations that are consistent with the conjunctive representations found in the ANN.

### Removing task context impairs task performance activity

We next sought to evaluate the importance of including task rule activations in model performance. To remove context activity, we lesioned all connections from the ENN’s rule input layer to the hidden layer. This was achieved by setting all resting-state FC connections from the context input layer to 0 (Fig. 7f). As hypothesized, model performance was at chance without task context activations (RH accuracy=50.00%, p=0.46; LH=50.00%, p=0.46; Fig. 7h,i). This demonstrated that the model implemented a representational transformation from task context and sensory stimulus activations to the correct motor response activations.

### The influence of specific functional network topography

We next evaluated whether the empirically-estimated connectivity topography was critical to successful task performance. This involved shuffling the connectivity weights within the context and stimulus input layers 1000 times (Fig. 7g). While we hypothesized that the specific resting-state FC topography would be critical to task performance, we found that shuffling connectivity patterns yielded a very high variance distribution of task performance (Fig. 7h). As expected, the mean across all connectivity shuffles were approximately at chance for both hands (RH mean accuracy=51.22%, p=0.44; LH mean accuracy=50.77%, p=0.47). However, we found that there were some connectivity configurations that would significantly improve task performance, and other connectivity configurations that would yield significant below chance task performance. Notably, the FC topography that was estimated from resting-state fMRI (the full S-R model, without shuffling; Fig. 7b) performed greater than 86.4% of all connectivity reconfigurations in RH responses, and greater than 97.0% of all connectivity reconfigurations for LH responses. This indicates that, among all possible connectivity patterns, the weights derived from resting-state fMRI stood out as especially effective at producing correct task performance.

## Discussion

Determining how the human brain transforms incoming stimuli into accurate task behavior would fill a critical gap in understanding how the brain implements cognitive processes^11, 33, 34^. To address this gap, we built a task-performing ENN from empirical data to identify the brain network mechanisms associated with representational transformations during a complex cognitive control task. This ENN was based on the conceptual frameworks provided by the Guided Activation and Flexible Hub theories, yet a lack of specificity regarding the functional equivalents of hypothesized components (e.g., the context and hidden layers) required us to perform additional theoretical and empirical work to find those equivalents. First, we identified brain vertices that were selective for task rules, sensory stimuli, motor responses, and conjunctions to be included as candidate areas to test the theories. Second, we mapped resting-state FC weights between these areas using multiple linear regression. Finally, using activity flow mapping, we found that incoming sensory and task rule activations were transformed via conjunction hubs to produce above-chance behavioral predictions of outgoing motor response activations. Thus, we not only identified *where* in the brain different task-related activations could be identified, but also *how* these task activations are transformed into behaviorally relevant motor activations through network computations supported by the brain’s intrinsic network organization.

Collectively, these findings suggest that flexible cognitive control is implemented by guided activations, as originally suggested by the Guided Activation Theory^3^. However, to more fully test this theory than prior work we had to substantially expand it, such as identifying the functional equivalents of hypothesized components in the human brain. Some of these components were not located where originally hypothesized. For instance, rather than context representations being confined to lateral prefrontal cortex, we found such representations distributed throughout the brain, with especially strong representation in the dorsal attention network (only a portion of which is in lateral prefrontal cortex). Similarly, the theory predicted the hidden layer (conjunction hubs) to be in non-prefrontal association cortex, yet we found some conjunction hubs in lateral prefrontal cortex and associated cognitive control networks (CCNs). Thus, we have empirically confirmed the broader Guided Activation Theory while expanding and refining it.

The present results build on the Flexible Hub theory and other findings emphasizing the role of CCNs in highly flexible cognition^1, 29, 35, 36^. Previous work on the Flexible Hub theory focused on characterizing rapid updates to task rule representations, finding that CCNs represent rules compositionally in both activity^7, 10, 36^ and FC^15, 16^ patterns. The present results build on those earlier findings, demonstrating that CCNs and other networks flexibly compose rule representations, since the ENN rule activation inputs contained three rules whose fMRI activity patterns were added compositionally to create the full task context. Critically, however, we found that these compositional codes were not enough to implement flexible task performance. Rather, conjunctive representations that conjoin rule and stimulus representations^9, 12^ were required to interact nonlinearly with these compositional representations. Interestingly, Kikumoto and Mayr recently demonstrated that conjunctive representations are critical to controlling motor responses, finding that the strength of conjunctive representations was associated with the success of motor responses^37^. Our results are consistent with those findings, showing that without conjunctive representations producing conditional interactions (i.e., through conjunction hub lesioning), the task performance of the ENN was substantially impaired. However, our results also differ from that study, since the conjunctive representations we identified were not simply multiplicative interactions between context and stimuli activations. Instead, our conjunctive representations are consistent with the biased competition theory of attention, where additive computations were passed through a nonlinearity^32^. It will be important for future work to distinguish the content of these two types of stimulus and rule conjunctions, and whether the representational content of conjunctions simultaneously contain both stimulus and rule content, or if instead rule and stimulus conjunctions collapse into contingency states for action selection as previously reported in computational models^38^. Nevertheless, these findings provide evidence to fill an important gap within the Flexible Hub framework, suggesting that the flexibility of rule updates are useful insofar as they can be integrated to form conjunctions with stimulus activity.

The ENN characterized the representational transformations required for task input activations to generate accurate output activations (in motor cortex) directly from data. Model parameters, such as unit identification and inter-unit connectivity estimation, were estimated *without optimizing for task performance*. This contrasts with mainstream machine learning techniques that iteratively train ANNs that directly optimize for behavior^13, 19, 20, 39, 40^. Our approach enabled the construction of functioning ENNs with above-chance task performance without optimizing for behavior; instead, we were able to derive parameters from empirical neural data alone. Theoretically, the results presented here are consistent with the goals of the Dynamic Causal Modeling (DCM) framework, which aim to identify the latent variables underlying input-output state transformations during tasks^41, 42^. However, in contrast to DCM, the present study 1) uses intrinsic rest FC to 2) build predictive models of task-evoked activity patterns coding for motor responses, which are then 3) tested against empirical activity patterns and task-appropriate behavior to assess model validity. These results suggest that the human brain’s intrinsic network architecture, as estimated with human fMRI data, is informative regarding the design of task-performing functioning models of cognitive computation.

We showed that the specific FC topography could predict inter-area transformations of fMRI activations. In contrast, shuffling these specific inter-area FC topographies yielded ENNs with highly variable task performances, suggesting the computational utility of the empirically-estimated FC patterns. Previous work has illustrated that the functional network architecture of the brain emerges from a structural backbone^43–48^. Building on this work, we recently proposed that the functional network architecture of the brain can be used to build network coding models – models of brain function that describe the encoding and decoding of task-relevant brain activity constrained by connectivity^49^. Related proposals have also been put forward in the electron microscopy connectomics literature, suggesting that structural wiring diagrams of the brain can inform functional models of biological systems (e.g., the drosophila’s visual system or the human brain’s intrinsic memory capacity)^47, 50, 51^. In addition, work in mean-field network models have revealed a direct link between connectivity and computations, finding that low-dimensional connectivity patterns (which also exist in fMRI data^52^) are useful for performing tasks^53^. Consistent with these proposals, our findings establish the feasibility of leveraging intrinsic FC to model representational transformations from sensory stimuli to motor responses during context-dependent tasks.

Despite the present study providing strong evidence that the estimated functional network model can perform tasks, several theoretical and methodological limitations remain. First, though we perform numerous control analyses by either lesioning or altering the ENN architecture (Fig. 7), the space of alternative possible models that can potentially achieve similar (if not better) task performances is large. For example, here we included only a single hidden layer (one layer of ‘conjunction hubs’). However, it is possible – if not probable – that such transformations actually involve a large sequence of transformations, similar to how the ventral visual stream transforms visual input into object codes, from V1 to inferior temporal cortex^20, 21^. Furthermore, recent work has suggested that conjunctive representations emerge at very fast timescales relative to the BOLD signal^9, 12^. However, the present study only focused on predicting conjunctive representations (using task context and stimulus activations as inputs to the ENN) in putative conjunction hubs (which did not require explicit estimation of conjunctions, but instead the representational relations between task context and stimulus activations). It is therefore likely that the identification of conjunctive representations is dependent on both specific task demands and the targeted level of analysis (e.g., neuronal circuits versus large-scale functional networks). Here we opted for the simplest possible network model that involved conjunction hubs at the level of large-scale functional networks. Starting from this simple model allowed us to reduce potential extraneous assumptions and model complexity (such as modeling the extraction of stimulus features from early visual areas and instead identifying late-stage sensory features, or cortical-subcortical interactions for action selection^54, 55^), which likely would have been necessary in more complex and detailed models. However, the current findings provide a strong foundation for future studies to unpack the mechanisms of finer-grained computations important for adaptive behavior.

Another assumption in the ENN was that activations were guided by additive connectivity weights. Additive connectivity weights assume inter-area predicted activations are the sum of source activations weighted by connections. One potential alternative (among others) would have been multiplicative guided activations; weighted activations that are multiplied (rather than summed) from incoming areas, which has been previously proposed as a potential alternative to designing ANNs^56^. However, several recent studies have suggested that inter-area activations are predicted via additive connectivity weights in both human fMRI^8, 18^, the primate visual system^22^, and the drosophila’s visual system^47^, suggesting that using additive connectivity weights is an appropriate model for how the brain implements computations. Nevertheless, it will be important for future work to directly adjudicate between potential alternatives (like multiplicative connectivity weights) in neural implementations of cognitive processes.

Finally, another limitation is that we constructed the ENN model without recurrent interactions, which are known to play a large role in neural computation^57, 58^. However, we still successfully captured some temporal dynamics, since activations from different task features (rule encoding and stimuli) were estimated from distinct temporal windows. We also note that though we estimate task encoding activity from the encoding period only, we modeled the result of persistent (potentially recurrent) activity in rule-representing regions by holding encoding activity in those regions constant across the delay period to the stimulus period. Nevertheless, though temporal dynamics (with recurrent feedback) likely play a role in shaping cognitive computations, we illustrate here that simple dynamics (i.e., rules + sensory inputs → conjunction hubs → motor outputs) involving the interplay of static activation patterns are sufficient to model representational transformations. We also tested the modeled cognitive transformations at the group level, limiting our ability to link individual task performance with individualized ENNs. However, our findings still significantly advance current understanding of how the brain transforms task context and stimulus activations into motor activations for behavior through computations implemented by intrinsic FC organization. Nevertheless, it will be important for future studies to construct individualized task-performing brain models that can simulate temporal and recurrent dynamics constrained by empirical data, as this can provide a more detailed computational account of the representational transformations that contribute to individual behavioral variability.

In conclusion, we constructed an ENN model from brain connectivity data that was capable of performing a complex cognitive control task. The model’s overall architecture was consistent with the prominent Guided Activation Theory, effectively validating the general form of that theory while substantially expanding it by revealing where and how its abstract functional components are implemented in the human brain. More broadly, this study illustrates an alternative perspective to the standard approach of using learning algorithms to train neural networks to perform tasks. Instead, brain data can be converted into generative neural network models that perform tasks, revealing how the brain generates that task performance. We expect that these findings will drive new investigations into the neural implementation of cognitive computations, providing dual insight into how the brain implements cognitive processes and how such knowledge can inform the design of ANN architectures.

## Methods

### Participants

Data were collected from 106 human participants across two different sessions (a behavioral and an imaging session). Participants were recruited from the Rutgers University-Newark community and neighboring communities. Technical error during MRI acquisition resulted in removing six participants from the study. Four additional participants were removed from the study because they did not complete the behavior-only session. fMRI analysis was performed on the remaining 96 participants (54 females). All participants gave informed consent according to the protocol approved by the Rutgers University Institutional Review Board. The average age of the participants that were included for analysis was 22.06, with a standard deviation of 3.84. Additional details regarding this participant cohort have been previously reported^59^.

### C-PRO task paradigm

We used the Concrete Permuted Operations (C-PRO) paradigm (Fig. 2a) during fMRI acquisition, and used a computationally analogous task when training our ANN model. The details of this task are described below, and are adapted from a previous study^8^.

The C-PRO paradigm is a modified version of the original PRO paradigm introduced in Cole et al., (2010)^60^. Briefly, the C-PRO cognitive paradigm permutes specific task rules from three different rule domains (logical decision, sensory semantic, and motor response) to generate dozens of novel and unique task contexts. This creates a context-rich dataset in the task configuration domain akin in some ways to movies and other condition-rich datasets used to investigate visual and auditory domains^5^. The primary modification of the C-PRO paradigm from the PRO paradigm was to use concrete, sensory (simultaneously presented visual and auditory) stimuli, as opposed to the abstract, linguistic stimuli in the original paradigm. Visual stimuli included either horizontally or vertically oriented bars with either blue or red coloring. Simultaneously presented auditory stimuli included continuous (constant) or non-continuous (non-constant, i.e., “beeping”) tones presented at high (3000Hz) or low (300Hz) frequencies. Fig. 2a demonstrates two example task-rule sets for “Task 1” and “Task 64”. The paradigm was presented using E-Prime software version 2.0.10.353^61^.

Each rule domain (logic, sensory, and motor) consisted of four specific rules, while each task context was a combination of one rule from each rule domain. A total of 64 unique task contexts (4 logic rules x 4 sensory rules x 4 motor rules) were possible, and each unique task set was presented twice for a total of 128 task miniblocks. Identical task sets were not presented in consecutive blocks. Each task miniblock included three trials, each consisting of two sequentially presented instances of simultaneous audiovisual stimuli. A task block began with a 3925 ms encoding screen (5 TRs), followed by a jittered delay ranging from 1570 ms to 6280 ms (2 – 8 TRs; randomly selected). Following the jittered delay, three trials were presented for 2355 ms (3 TRs), each with an inter-trial interval of 1570 ms (2 TRs). A second jittered delay followed the third trial, lasting 7850 ms to 12560 ms (10-16 TRs; randomly selected). A task block lasted a total of 28260 ms (36 TRs). Subjects were trained on four of the 64 task contexts for 30 minutes prior to the fMRI session. The four practiced rule sets were selected such that all 12 rules were equally practiced. There were 16 such groups of four task sets possible, and the task sets chosen to be practiced were counterbalanced across subjects. Subjects’ mean performance across all trials performed in the scanner was 84% (median=86%) with a standard deviation of 9% (min=51%; max=96%). All subjects performed statistically above chance (25%).

### fMRI acquisition and preprocessing

The following fMRI acquisition details is taken from a previous study that used the identical protocol (and a subset of the data)^8^.

Data were collected at the Rutgers University Brain Imaging Center (RUBIC). Whole-brain multiband echo-planar imaging (EPI) acquisitions were collected with a 32-channel head coil on a 3T Siemens Trio MRI scanner with TR=785 ms, TE=34.8 ms, flip angle=55°, Bandwidth 1924/Hz/Px, in-plane FoV read=208 mm, 72 slices, 2.0 mm isotropic voxels, with a multiband acceleration factor of 8. Whole-brain high-resolution T1-weighted and T2-weighted anatomical scans were also collected with 0.8 mm isotropic voxels. Spin echo field maps were collected in both the anterior to posterior direction and the posterior to anterior direction in accordance with the Human Connectome Project preprocessing pipeline^62^. A resting-state scan was collected for 14 minutes (1070 TRs), prior to the task scans. Eight task scans were subsequently collected, each spanning 7 minutes and 36 seconds (581 TRs). Each of the eight task runs (in addition to all other MRI data) were collected consecutively with short breaks in between (subjects did not leave the scanner).

### fMRI Preprocessing

The following details are adapted from a previous study that used the same preprocessing scheme on a different data set^63^.

Resting-state and task-state fMRI data were minimally preprocessed using the publicly available Human Connectome Project minimal preprocessing pipeline version 3.5.0. This pipeline included anatomical reconstruction and segmentation, EPI reconstruction, segmentation, spatial normalization to standard template, intensity normalization, and motion correction^64^. After minimal preprocessing, additional custom preprocessing was conducted on CIFTI 64k grayordinate standard space for vertex-wise analyses using a surface based atlas^25^. This included removal of the first five frames of each run, de-meaning and de-trending the time series, and performing nuisance regression on the minimally preprocessed data^64^. We removed motion parameters and physiological noise during nuisance regression. This included six motion parameters, their derivatives, and the quadratics of those parameters (24 motion regressors in total). We applied aCompCor on the physiological time series extracted from the white matter and ventricle voxels (5 components each extracted volumetrically)^65^. We additionally included the derivatives of each component time series, and the quadratics of the original and derivative time series (40 physiological noise regressors in total). This combination of motion and physiological noise regressors totaled 64 nuisance parameters, and is a variant of previously benchmarked nuisance regression models^64^.

### fMRI task activation estimation

We performed a standard task GLM analysis on fMRI task data to estimate task-evoked activations from different conditions. Task GLMs were fit for each subject separately, but using the fully concatenated task data set (concatenated across 8 runs). We obtained regressors for each task rule (during the encoding period), each stimulus pair combination (during stimulus presentation), and each motor response (during button presses). For task rules, we obtained 12 regressors that were fit during the encoding period, which lasted 3925ms (5 TRs). For logic rules, we obtained regressors for “both”, “not both”, “either”, and “neither” rules. For sensory rules, we obtained regressors for “red”, “vertical”, “high”, and “constant” rules. For motor rules, we obtained regressors for “left middle”, “left index”, “right middle”, and “right index” rules. Note that a given encoding period contained overlapping regressors from each of the logic, sensory, and motor rule domains. However, the regressors were not collinear since specific rule instances were counterbalanced across all encoding blocks.

To obtain activations for sensory stimuli, we fit regressors for each stimulus pair. For example, for the color dimensions of a stimulus, we fit separate regressors for the presentation of red-red, red-blue, blue-red, and blue-blue stimulus pairs. This was done (rather than fitting regressors for just red or blue) due to the inability to temporally separate individual stimuli with fMRI’s low sampling rate. Thus, there were 16 stimulus regressors (four conditions for each stimulus dimension: color, orientation, pitch, continuity). Stimulus pairs were presented after a delay period, and lasted 2355ms (3 TRs). Note that a given stimulus presentation period contained overlapping regressors from four different conditions, one from each stimulus dimension. However, the stimulus regressors were not collinear since stimulus pairings were counterbalanced across all stimulus presentation periods (e.g., red-red stimuli were not exclusively presented with vertical-vertical stimuli).

Finally, to obtain activations for motor responses (finger button presses), we fit a regressor for each motor response. There were four regressors for motor responses, one for each finger (i.e., left middle, left index, right middle, right index fingers). Responses overlapped with the stimulus period, so we fit regressors for each button press during the 2355ms (3 TR) window during stimulus presentations. Note, however, that while response regressors overlapped with stimulus regressors, estimated response activations were independent from stimulus activations. There were two reasons for this: 1) Motor response and stimulus regressors were equally independent from each other due to counterbalancing across conditions (e.g., the same stimulus elicited all other motor responses equally; see Supplementary Fig. 9); 2) Motor response and stimulus activations were estimated in the same task GLM model (multiple linear regression, across the counterbalanced conditions), such that activations associated with each condition contained unique variance. (This is because multiple linear regression conditions on all other regressors.) A strong validation of this approach was that the finger activations could be reliably extracted according to the appropriate topographic organization in somatomotor cortex (Fig. 4c).

For a schematic of how task GLMs were performed, see Supplementary Fig. 8. For the task design matrix of an example subject, see Supplementary Fig. 9.

### fMRI decoding: Identifying sensory stimulus activations

Decoding analyses were performed to identify the brain areas that contained relevant task context and sensory stimulus activations. To identify the brain areas that contained relevant sensory stimulus representation, we performed four, four-way decoding analyses on each stimulus dimension: color (vision), orientation (vision), pitch (audition), constant (audition). For color stimulus representations, we decoded activation patterns where the stimulus pairs were red-red, red-blue, blue-red, and blue-blue. For orientation stimulus representations, we decoded activation patterns where the stimulus pairs were vertical-vertical, vertical-horizontal, horizontal-vertical, horizontal-horizontal. For pitch stimulus representations, we decoded activation patterns where the stimulus pairs were high-high, high-low, low-high, and low-low. Finally, for constant (beeping) stimulus representations, we decoded activation patterns where the stimulus pairs were constant-constant, constant-beeping, beeping-constant, beeping-beeping.

Decoding analyses were performed using the vertices within each parcel as decoding features. We limited decoding to visual network parcels for decoding visual stimulus features, and auditory network parcels for decoding auditory stimulus features. Visual parcels were defined as the VIS1 and VIS2 networks in Ji et al. (2019)^26^, and auditory networks as the AUD network. We performed a group-level decoding analysis with a 12-fold cross-validation scheme. We used a minimum-distance/nearest-neighbor classifier (based on Pearson’s correlation score), where a test set sample would be classified as the condition whose centroid is closest to in the activation pattern space^24^. P-values were calculated using a binomial test. Statistical significance was assessed using a false discovery rate (FDR) corrected threshold of p<0.05 across all 360 regions. To ensure robustness of all fMRI decoding analyses, we additionally performed logistic classifications (linear decoding) to compare with minimum-distance-based classifiers. (See also refs^66, 67^ for comparing distance versus linear-based similarity measures.) In general, there were no differences between the two decoding schemes, although in one instance (task-rule decoding), minimum-distance classifiers significantly outperformed logistic classification (Supplementary Fig. 7)

### fMRI decoding: Identifying task rule activations

To identify the brain areas that contained task rule activations, we performed a 12-way decoding analysis on the activation patterns for each of the 12 task rules. We used the same decoding and cross-validation scheme as above (for identifying sensory stimulus representations). However, we ran the decoding analyses on all 360 parcels, given previous evidence that task rule activity is widely distributed across cortex^8^. P-values were calculated using a binomial test. Statistical significance was assessed using an FDR-corrected threshold of p<0.05 across all 360 regions.

### fMRI activation analysis: Identifying motor response activations

To identify the brain areas/vertices that contained motor response activity, we performed univariate analyses to identify the finger press activations in motor cortex. In contrast to identifying other task components via decoding analyses (e.g., rules and stimuli), we were able to use simpler univariate methods (i.e., t-tests) to identify motor response vertices. This was because the identification of index versus middle finger response vertices did not require a multi-way decoding analysis (unlike stimulus and rule conditions, which had 4 and 12 conditions, respectively). Instead, motor response identification only required identifying vertex-wise receptive field activations corresponding to each finger (suitable for a 2-way univariate test). This provided a more constrained and biologically interpretable receptive field for each response activation, rather than including the entire primary cortex.

We performed two univariate activation contrasts, identifying index and middle finger activations on each hand. For each hand, we performed a two-sided group paired (by subject) t-test contrasting index versus middle finger activations. We constrained our analyses to include only vertices in the somatomotor network. Statistical significance was assessed using an FDR-corrected p<0.05 threshold, resulting in a set of vertices that were selective to button press representations in motor cortex (see Fig. 4c).

We subsequently performed a cross-validated decoding analysis on vertices within the motor cortex to establish a baseline noise ceiling of motor response decodability (see Fig. 7a,h). We decoded finger presses on each hand separately. To identify specific vertices for selective response conditions, we performed feature selection on each cross-validation loop separately to avoid circularity. Feature selection criteria (within each cross-validation loop) were vertices that survived a p<0.05 threshold (using a paired t-test). We performed a 4-fold cross validation scheme using a minimum-distance classifier, bootstrapping training samples for each fold. Moreover, because the decoding analysis was limited to a single ROI (as opposed to across many parcels/ROIs), we were able to compute confidence intervals (by performing multiple cross-validation folds) and run nonparametric permutation tests since it was computationally tractable. We ran each cross-validation scheme 1000 times to generate confidence intervals. Null distributions were computed by randomly permuting labels 1000 times. P-values were computed by comparing the null distribution against each of the bootstrapped accuracy values, then averaging across p-values.

### Identifying conjunctive representations: ANN construction

We trained a simple feedforward ANN (with layers organized according to the Guided Activation Theory) on a computationally analogous form of the C-PRO task. This enabled us to investigate how task rule and stimulus activations conjoin into conjunctive activations in an ANN’s hidden layer.

To model the task context input layer, we designated an input unit for each task rule across all rule domains. Thus, we had 12 units in the task context layer. A specific task context (or rule set) would selectively activate three of the 12 units; one logic rule, one sensory rule, and one motor rule. Input activations were either 0 or 1, indicating an active or inactive state.

To model the stimulus input layer, we designated an input unit for a stimulus pair for each sensory dimension. To isolate visual color stimulus pairings, we designated input units for a red-red pairing, red-blue pairing, blue-red pairing, and blue-blue pairing. (Note that each unit represented a stimulus pair because the ANN had no temporal dynamics to present consecutive stimuli.) To isolate visual orientation stimulus pairings, we designated inputs for a vertical-vertical, vertical-horizontal, horizontal-vertical, and horizontal-horizontal stimulus pairing. To isolate auditory pitch stimulus pairings, we designated input units for high-high, high-low, low-high, and low-low frequency combinations. Finally, to isolate auditory continuity stimulus pairings (i.e., whether an auditory tone was constant or beeping), we designated input units for constant-constant, constant-beeping, beeping-constant, and beeping-beeping. Altogether, across the four sensory domains, we obtained 16 different sensory stimulus pair input units. For a given trial, four units would be activated to simulate a sensory stimulus combination (one unit per sensory domain). For example, a single trial might observe red-red (color), vertical-horizontal (orientation), low-high (pitch), constant-beeping (continuity) stimulus combination. Thus, to simulate an entire trial including both context and sensory stimuli, 7/28 possible input units would be activated.

We constructed our ANN with two hidden layers containing 1280 units each. This choice was due to recent counterintuitive evidence suggesting that the learning dynamics of extremely high-dimensional ANNs (i.e., those with many network parameters to tune) naturally protect against overfitting, supporting generalized solutions^68^. Moreover, we found that across many initializations, the representational geometry identified in the ANN’s hidden layer was highly replicable. Finally, our output layer contained four units, one for each motor response (corresponding to left middle, left index, right middle, right index finger presses).

The ANN transformed a 28-element input vector (representing a specific trial instance) into a 4-element response vector, and obeyed the equation

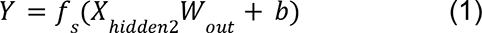

where *Y* corresponds to the 4-element response vector, *f_s_* is a sigmoid function, *W_out_* corresponds to the connectivity weight matrix between the hidden and output layer, *b* is a bias term, and *X_hidden2_* is the activity vector of the 2nd hidden layer. *X_hidden2_* was obtained by the equation

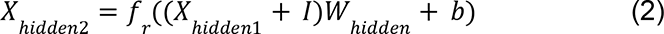

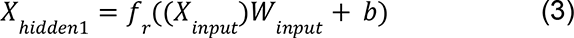

Where *f_r_* is a rectified linear function (ReLU), *W_hidden_* is the connectivity matrix between the hidden layers, *X_hidden1_* corresponds to the 1st hidden layer activations that contain trial information, *X_input_* is the input layer, *W_input_* is the connectivity matrix between the input and 1st hidden layer, and *I*is a noise vector sampled from a normal distribution with 0-mean and 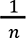-variance, where *n* refers to the number of hidden units.

### Identifying conjunctive representations: ANN training

The ANN was trained by minimizing the mean squared error between the network’s outputs and the correct target output. The mean squared error was computed using a mini-batch approach, where each mini-batch consisted of 192 distinct trials. (Each of the 64 unique task contexts were presented three times (with randomly sampled stimuli) in each mini-batch. Training was optimized using Adam, a variant of stochastic gradient descent^69^. We used the default parameters in PyTorch (version 1.0.1), with a learning rate of 0.0001. Training was stopped when the last 1000 mini-batches achieved over 99.5% average accuracy on the task. This performance was achieved after roughly 10,000 mini-batches (or 1,920,000 trials). Weights and biases were initialized with a uniform distribution 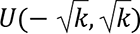, where 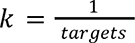, where ‘targets’ represents the number of units in the next layer. We note that the representational geometry we observed in the hidden layer was robust to different initializations and hyperparameter choices, indicating strong test-retest reliability of learned hidden layer representations (Supplementary Fig. 6). For example, the ANN’s hidden layer RSM was also consistent across different ANN instantiations with different hidden layer sizes (Supplementary Fig. 6). We also ran an additional null model in which we randomly shuffled trial labels during training, arbitrarily remapping rule- and stimulus-response mappings. We found that with the ANN architecture and parameters, the ANN could not learn the task with shuffled labels since the hierarchical reasoning structure of the C-PRO task was destroyed with shuffling. This suggested that the unshuffled ANN we used did not learn the C-PRO task with a memorization strategy.

We note that the ANN is entirely distinct from the ENN, and that only the ANN used gradient descent for training. The sole purpose of the ANN was to identify conjunctive representations in the ANN’s hidden layer, which was in turn used to identify conjunctive representations in empirical data (through matching the representational similarity matrices of the ANN and empirical data described below).

### Identifying conjunctive representations: ANN representational analysis

We extracted the representational geometry of the ANN’s 2nd hidden layer using representational similarity analysis (RSA)^70^. This was done to understand how task rule and stimulus activations were transformed in the hidden layer. To extract the representational geometry of the hidden layer, we systematically activated a single unit in the input layer (which corresponded to either a task rule or sensory stimulus pair), and estimated the corresponding hidden layer activations (using trained connectivity weights). This resulted in a total of 28 (12 task rules and 16 sensory stimuli combinations) activation patterns. The representational similarity matrix (RSM) was obtained by computing the Pearson correlation between the hidden layer activation patterns for all 28 conditions.

Identification of the control ANN’s hidden layer RSM (Supplementary Fig. 4b) was obtained by randomly shuffling all weights and biases (within each layer) after training. This preserved the distribution of the weights and biases of the trained ANN, while impairing the ANN’s ability to properly perform the task. Shuffling was performed 10,000 times, and the null RSM was obtained by averaging the RSMs across permutations.

### Identifying conjunctive representations: fMRI analysis

We compared the representational geometry of the ANN’s hidden layer to the representational geometry of each brain parcel. This was possible because we extracted the exact same set of activation patterns (e.g., activations for task rules and sensory stimuli) in empirical data as our ANN model, enabling a direct comparison of representations. The representational geometry was estimated as the representational similarity matrix (RSM) of all task rules and sensory stimuli conditions.

We first estimated the empirical RSMs for every brain parcel separately in the Glasser et al. (2016) atlas. This was done by comparing the activation patterns of each of the 28 task conditions using the vertices within each parcel (12 task rule activations, 16 sensory stimulus activations). We then applied a Fisher’s *z*-transform on both the empirical and ANN’s RSMs, and then estimated the Spearman’s rank correlation between the Fisher’s *z*-transformed ANN and empirical RSMs (using the upper triangle values only). This procedure was performed on the RSM of every brain parcel, providing a similarity score between each brain parcel’s and the ANN’s representational geometry. For our main analysis, we selected the top 10 parcels with highest similarity to the ANN’s hidden layer. However, we also performed additional analyses using the top 20, 30, and 40 parcels.

### FC weight estimation

We estimated resting-state FC to identify weights between layers in our empirical model. This was similar to our previously published approach that identified FC weights between pairs of brain regions^8^. This involved identifying FC weight mappings between the task rule input layer to the hidden layer, sensory stimulus input layer to the hidden layer, and the hidden layer to the motor output layer. For each FC mapping, we estimated the vertex-to-vertex FC weights using principal components linear regression. Consistent with our prior studies^8, 18^, we used principal components regression because most layers had more vertices (i.e., predictors) than samples in our resting-state data (resting-state fMRI data contained 1065 TRs). Principal components regression first identifies a set of principal components from all of the vertex time series of the source layer (via principal component analysis), then fits those latent components to each target layer vertex time series using multiple linear regression. For all FC estimations, we used principal components regression with 500 components, as we have in prior work^8, 18^. Specifically, FC weights were estimated by fitting principal components to the regression equation

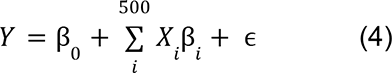

where *Y* corresponds to the t x n matrix with t time points and n vertices (i.e., the target vertices to be predicted), β_0_corresponds to a constant term, β*_i_* corresponds to the 1 x n matrix reflecting the mapping from the component time series onto the n target vertices, *X_i_* corresponds to the t x 1 component time series for component i, and ɛ corresponds to the error in the regression model. Note that *X*corresponds to the t x 500 component matrix obtained from a principal component analysis on the resting-state data from the source layer. Also note that these loadings onto these 500 components are saved for later, when task activation patterns from a source layer are projected onto a target layer. The loadings project the original vertex-wise task activation patterns in the source layer onto a lower-dimensional space enabling faster computations. A similar approach was used in a previous study ^71^. FC weights were computed for each individual separately, but then averaged across subjects to obtain group FC weights.

Note that in some cases, it was possible for overlap between the source and target vertices. (For example, some conjunction hub vertices may have coincided with the same vertices in the context layer.) In these cases, these overlapping vertices were excluded in the set of predictors (i.e., removed from the source layer) in the regression model.

### Simulating sensorimotor transformations with multi-step activity flow mapping

We generated predictions of motor response activations (in motor cortex) by assessing the correct motor response given a specific task context and sensory stimulus activation pattern (for additional details see Supplementary Fig. 1). For each subject, we simulated 960 pseudo-trials. (“Pseudo-trials” refer to simulated trials using estimated activations rather than the actual experimental trials subjects performed.) This consisted of the 64 unique task contexts each paired with 15 randomly sampled stimulus combinations for a total of 960 pseudo-trials. For a pseudo-trial, the task context input activation pattern was obtained by extracting the activation vector for the logic, sensory, and motor rule, and computing the mean rule vector (i.e., additive compositionality). The sensory stimulus input activation pattern was obtained by extracting the relevant sensory stimulus activation pattern. (Note that for a given pseudo-trials, we only extracted the activation pattern for the sensory feature of interest. For example, if the rule was “Red”, only color activation patterns would be extracted, and all other stimulus activations would be set to 0.) Thus, the context and sensory stimulus activation patterns could be defined as

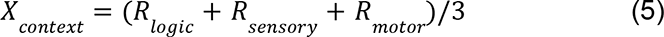

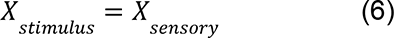

where *X*_*context*_ corresponds to the input activation pattern for task context, *R_logic_* corresponds to extracted logic rule activation pattern (e.g., “Both”, “Not Both”, “Either”, or “Neither”) obtained from the task GLM, *R_sensory_* corresponds to the extracted sensory rule activation pattern from the task GLM, *R*_*motor*_ corresponds to the extracted motor rule activation pattern from the task GLM, and *X_stimulus_* corresponds to the extracted sensory stimulus activation pattern that is indicated by the task context.

*X_context_* and *X_stimulus_* reflect the input activation patterns that were used to generate/predict motor response conditions. Importantly, these input activation patterns contained no information about the motor response, as illustrated by alternative null models (Fig. 7).

We used the FC weight maps to project *X_context_* and *X_stimulus_* onto the hidden/conjunction layer vertices. The projections (or predicted activation patterns on the hidden layer) were then thresholded to remove any negative BOLD predictions (i.e., values below the between-task-block resting baseline). This thresholding was used because it is equivalent to a rectified linear unit (ReLU), a commonly used nonlinear function in artificial neural networks^40^. Thus, the hidden layer was defined by

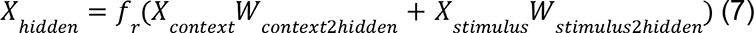

where *X_hidden_* corresponds to the predicted hidden layer activation pattern, *f_r_* is a ReLU function (i.e., *f_r_* (*x*) = *max*(*x*, 0)), *W_context2hidden_* corresponds to the resting-state FC weights between the context and hidden layer, and *W_stimulus2hidden_* corresponds to the resting-state FC weights between the stimulus and hidden layer. Note that all FC weights (*W_x_*) were computed using a principal component regression with 500 components. This requires that the vertex-wise activation space (e.g., *X_context_*) be projected onto component space such that we define

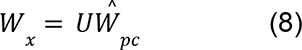

where *U* corresponds a *m* x 500 matrix which maps the source layer’s *m* vertices into component space, and *Ŵ*_*pc*_ is a 500 x *n* matrix that maps the components onto the target layer’s *n* vertices. (Note that *Ŵ*_*pc*_ corresponds to the regression coefficients from equation 4., and that both *U* and *Ŵ*_*pc*_ are estimated from resting-state data.) Thus, *W_x_* is an *m* x *n* transformation from a source layer’s spatial pattern to a target layer’s spatial pattern that is achieved through principal component regression on resting-state fMRI data.

Finally, we generated a predicted motor output response by computing

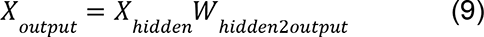

where *X_output_* corresponds to the predicted motor response (in motor cortex), and *W_hidden2output_* corresponds to the resting-state FC weights between the hidden and output layer. The full model computation can thus be formalized as

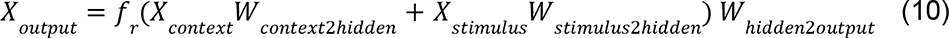

*X_output_* only yields a predicted activation pattern for the motor cortex for a given context and stimulus input activation pattern. To evaluate whether *X_output_* could successfully predict the *correct* motor response for a given pseudo-trials, we constructed an ideal ‘task solver’ that would indicate the correct motor response on a given pseudo-trial (Supplementary Fig. 1). This solver would then be used to identify the correct motor response activation pattern such that we could compare the predicted motor cortex activation with the actual motor cortex activation pattern.

We simulated 960 pseudo-trials per subject, randomly sampling context and stimulus input activation patterns. Because we sampled across the 64 task contexts equally (15 pseudo-trials per context), the correct motor responses were equally counterbalanced across 960 pseudo-trials. Thus, of the 960 simulated pseudo-trials for each subject, 240 pseudo-trials yielded a left middle, left index, right middle, and right index response each. For each of the 240 predicted motor response patterns we subsequently averaged across pseudo-trials such that we obtained 4 predicted motor response patterns for each subject. This choice was made for computational tractability (and boosting of signal-to-noise ratio), enabling us to downsample the large number of simulated pseudo-trials into predicted prototypical response activations for individual subjects. This reduced the number of samples the classifier had to train on 240-fold. Statistical evaluation of the 4 predicted (averaged) motor responses per subject was performed at the group level using a cross-validation scheme described below. See Supplementary Fig. 1b for a description of subject-versus group-level contributions to the ENN model.

### Statistical and permutation testing of predicted motor response activations

The simulated empirical model generated predicted activations of motor activations in motor cortex. However, the predictions would only be interesting if they resembled actual motor response activations directly estimated from the response period via task GLM. In other words, without a ground truth reference to the actual motor response activation patterns, the predicted activation patterns would hold little meaning. The simulated empirical model generated four predicted activation patterns corresponding to predicted motor responses for each subject. We also had four *actual* activation patterns corresponding to motor responses that were extracted from the motor response period using a standard task GLM for each subject. To test whether the predicted activation patterns conformed to the actual motor response activation patterns, we trained a decoder on the predicted motor response activations and tested on the actual motor response activations of held-out subjects. We used a 4-fold cross-validation decoding scheme (with a minimum-distance/Pearson correlation classifier), where training was performed on predicted motor activation patterns of 72 subjects, while testing was performed on the actual motor activation patterns of 24 held-out subjects. Training samples were randomly sampled with replacement. Training a decoder on the predicted activations and decoding the actual activations made this analysis consistent with a prediction perspective – we could test if, in the absence of any motor task activation, the ENN could predict actual motor response activation patterns that correspond to behavior.

Statistical significance was assessed using permutation tests. We permuted the labels of the predicted motor responses while testing on the actual motor responses. Null distributions are visualized in gray (Fig. 7h). For each model, we repeated the 4-fold cross-validation analysis 1000 times with correct labels to evaluate the robustness of the decoding accuracies. Statistical significance was assessed by generating a non-parametric p-value estimated from the null distribution for each iteration’s accuracy. Reported p-values were the average across all p-values for each model. Statistical significance was defined by a p<0.05 threshold.

### Data availability

All data related to this study are publicly available on OpenNeuro (https://openneuro.org/datasets/ds003701).

### Code availability

All code related to this study is publicly available on GitHub (https://github.com/ito-takuya/sr_enn).

## Acknowledgements

This project was supported by the US National Institutes of Health, under awards K99-R00 MH096901 and R01 MH109520. This material was also supported by the National Science Foundation under Grant No. BCS-1828528. The authors acknowledge the Office of Advanced Research Computing (OARC) at Rutgers, The State University of New Jersey for providing access to the Amarel cluster and associated research computing resources. The content is solely the responsibility of the authors and does not necessarily represent the official views of any of the funding agencies.

## Supplementary Figures

**Supplementary Figure 1.**
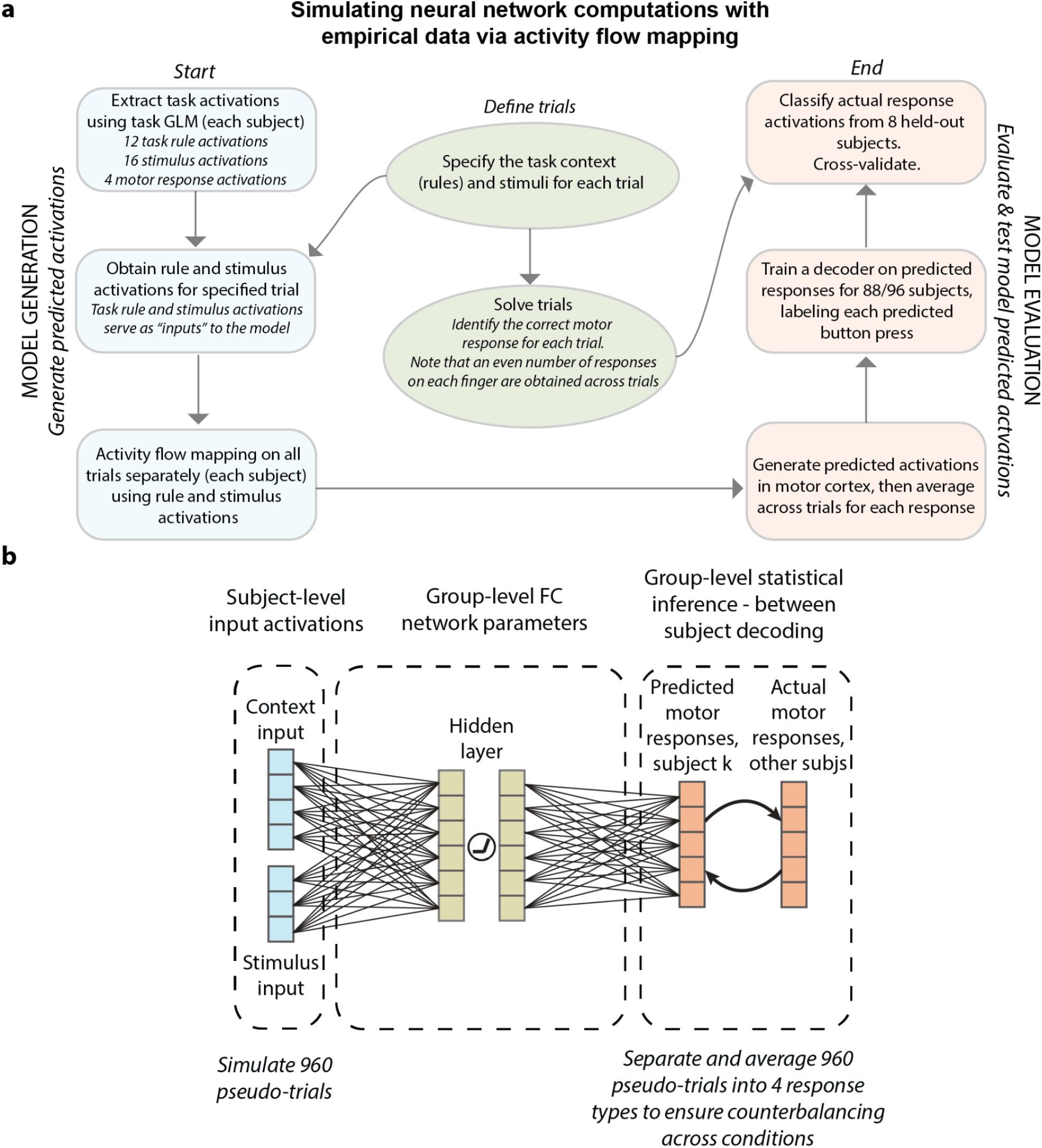
Additional details describing the ENN model and simulations. **a)** Flow chart describing neural network simulations with empirical data via activity flow mapping. We generate a subject’s predicted motor response activations using only task rule and sensory stimulus activation patterns as inputs. We then test these predictions against the actual motor response activations of other subjects. **b)** Detailing subject-specific versus group-level contributions of the ENN. The ENN produced group-level inferences on transforming rule and stimulus activations into motor response activations. Group-level inferences were assessed by between-subject decoding, training a model on predicted motor responses and predicted the actual motor response activations of *other subjects* in a cross-validated fashion. The core network parameters (e.g., resting-state FC weights) were estimated at the group-level. Input activations were estimated at the subject level to ensure that group-level predictions could be performed on predictions.

**Supplementary Figure 2.**
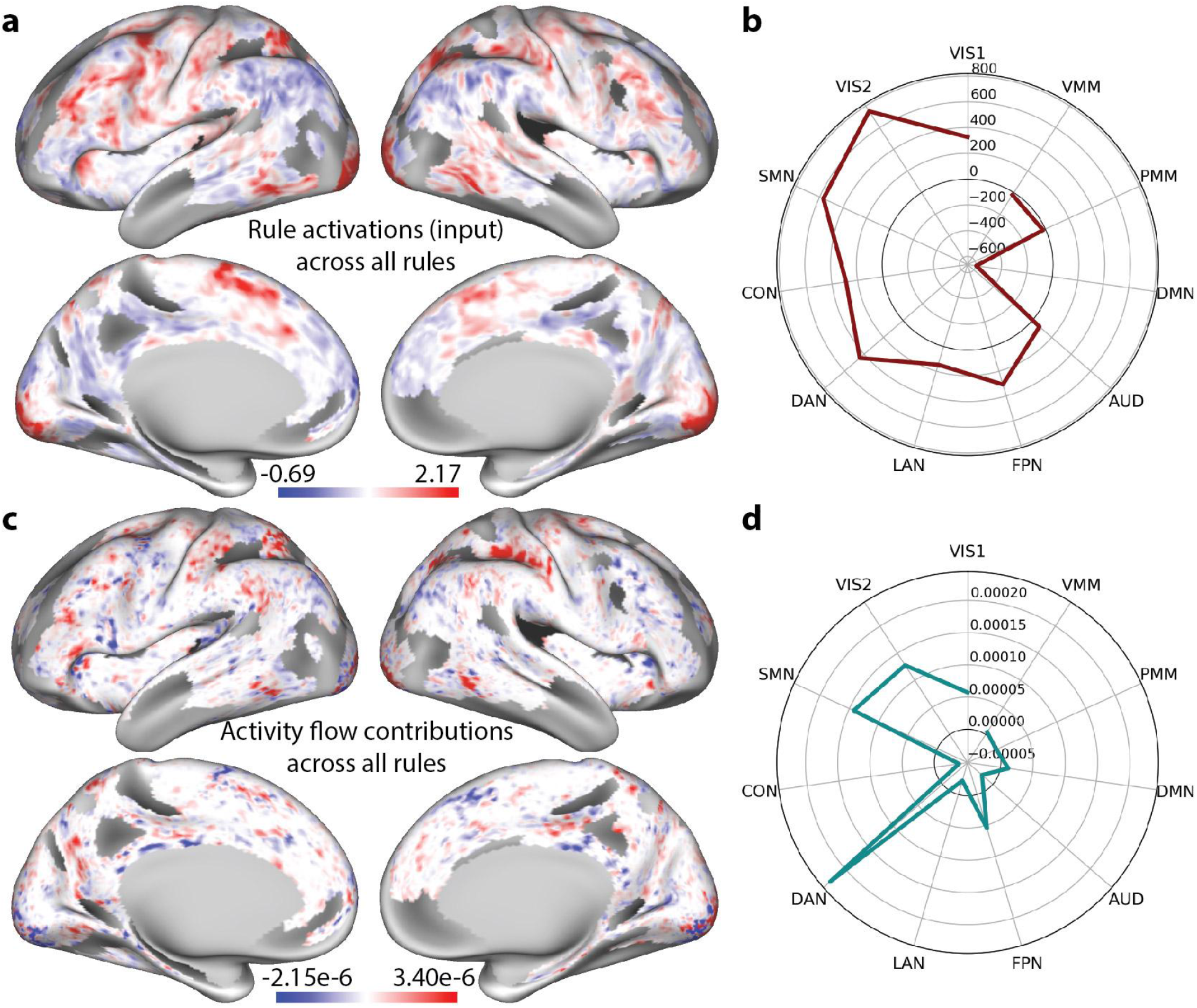
Comparison of task rule activations (in task rule regions) and their activity flow contributions (activation multiplied by FC weights) onto conjunction hubs. **a)** The task rule activations for every vertex in the rule regions, averaged across all rule sets. This map illustrates the activations that are fed in as inputs into the ENN. Units are in task GLM beta coefficients. **b)** The distribution of task rule activations, summed across all vertices within each functional network. Both sensorimotor and association networks appear to have strong rule-related activations. **c)** The activity flow contributions (task rule activations weighted by their FC weights with conjunction hubs) of each vertex in the task rule regions. The activity flow map is noticeably different from a), given that this visualization takes into consideration both the task rule activity and their FC with conjunction hubs. This map is obtained by averaging across the activity flow contributions to all vertices in conjunction hubs. **d)** The distribution of task rule activity flow contributions to conjunction regions, summed across all vertices within each functional network. Despite having large activations in sensorimotor regions in the original task rule activation map, the activity flow contributions from these networks are dampened by their FC to conjunction hubs. The DAN contributes the most activity flow to conjunction hubs. Network abbreviations: Primary visual (VIS1), Secondary visual (VIS2), Somatomotor (SMN), Cingulo-opercular (CON), Dorsal attention (DAN), Language (LAN), Frontoparietal (FPN), Auditory (AUD), Default Mode (DMN), Posterior multi-modal (PMM), Ventral multi-modal (VMM).

**Supplementary Figure 3.**
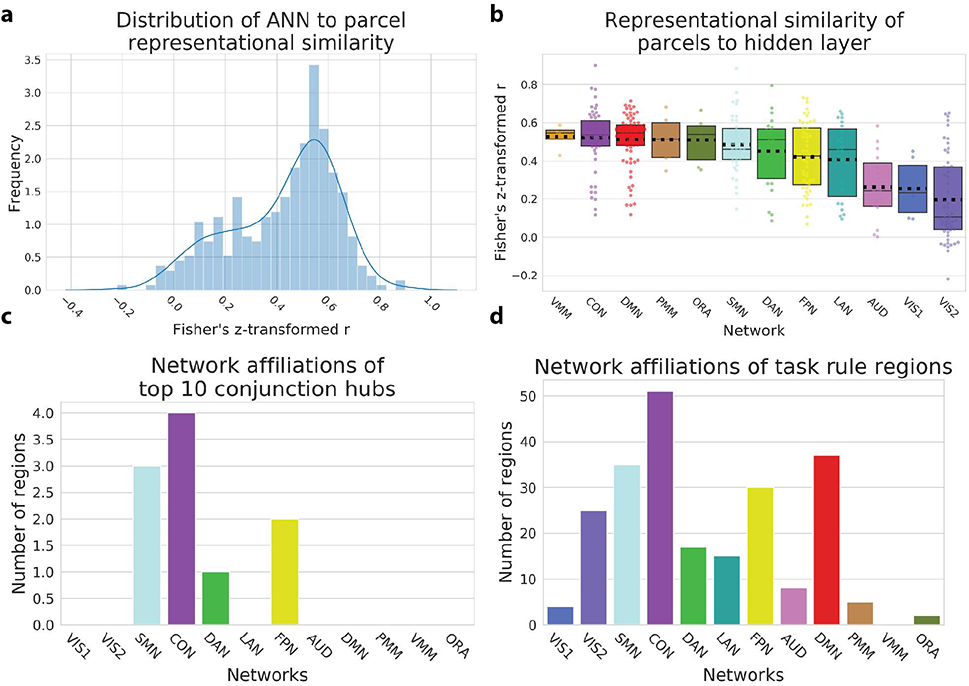
Distribution and network associations of conjunction hubs and the task rule input layer using a previously defined multimodal atlas and network partition^25, 26^. **a)** Histogram of the similarity (rank correlation) of the representational similarity matrix of the ANN’s hidden layer and each cortical parcel (Mean of distribution=0.43). **b)** Distribution of similarity scores for each functional network (each point reflects a different parcel belonging to the functional network). Boxplots reflect the interquartiles of the distribution, black dotted line the mean, and solid line the median. Strip plots reflect the entire distribution. **c)** The network affiliations of the 10 conjunction hub brain areas. **d)** Network affiliations of the 228 brain regions that contained decodable task rule activations. Network abbreviations: Primary visual (VIS1), Secondary visual (VIS2), Somatomotor (SMN), Cingulo-opercular (CON), Dorsal attention (DAN), Language (LAN), Frontoparietal (FPN), Auditory (AUD), Default Mode (DMN), Posterior multi-modal (PMM), Ventral multi-modal (VMM), Orbito-affective (ORN).

**Supplementary Figure 4.**
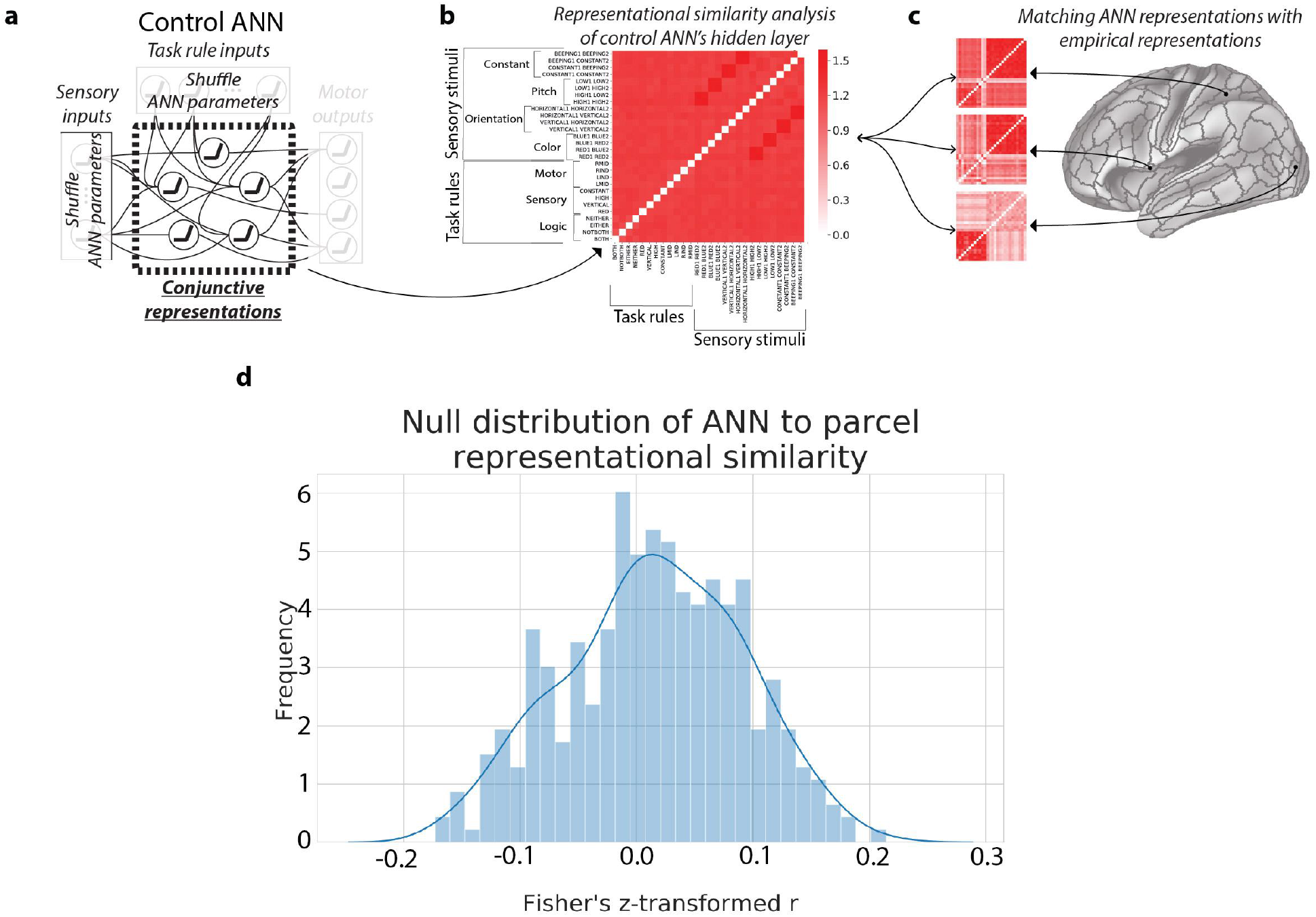
Evaluating the representational similarity of each brain parcel with a control ANN with shuffled parameters (e.g., weights). **a)** We generated 10,000 control ANNs, where after training, we randomly shuffled the parameters (i.e., weights and biases) within each layer. This was an appropriate control since this impaired the ANN’s ability to perform the task (at chance), while preserving the distribution of parameters within each layer. **b)** Using the control ANNs, we computed the averaged representational similarity matrix of the hidden layer. We found that the control ANN effectively obliterated any representational structure (dissimilarities) that were present in the unshuffled ANN (see Fig. 5b). **c)** As in Figure 5, we measured the similarity of the control ANN’s RSM with the RSMs of each brain parcel. **d)** Histogram of the similarity (rank correlation) of the RSM of the control ANN’s hidden layer and the RSMs of each cortical parcel (Mean of distribution=0.02). In contrast to the non-shuffled ANN’s hidden layer (see Supplementary Fig. 3a), the mean similarity (across all parcels) was near 0, suggesting that the correspondence between the ANN’s true hidden layer and the empirical data were highly non-trivial.

**Supplementary Figure 5.**
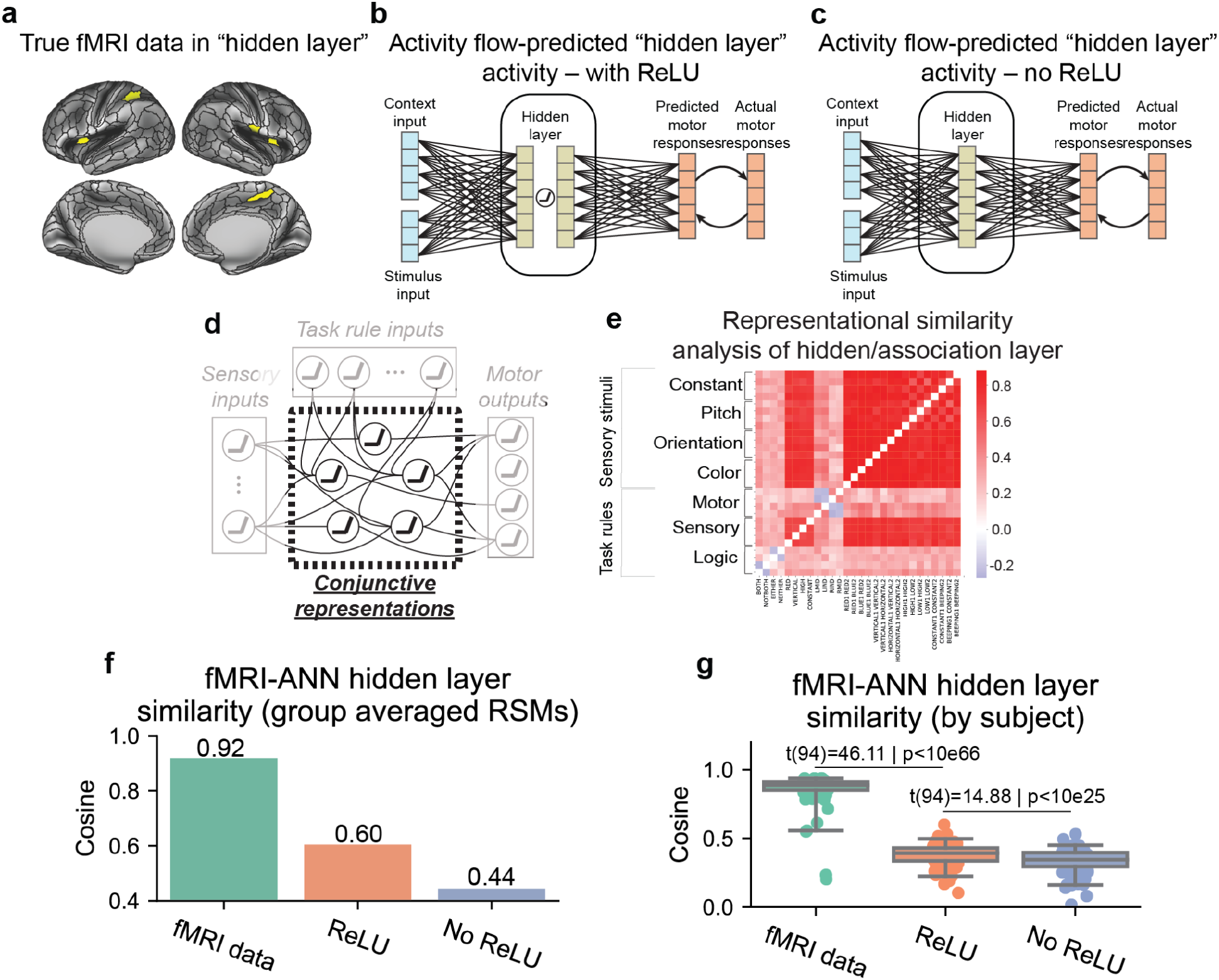
Inclusion of the ReLU in the hidden layer increased the similarity of activity flow-predicted hidden representations with the conjunctive hidden representations in the ANN. We calculated the ANN’s hidden layer representations with the **a)** empirical representations in the true conjunctive “hidden” layers of the fMRI data, **b)** the activity flow-predicted hidden representations in those same parcels after the ReLU was applied, and **c)** the activity flow-predicted hidden representations in those same parcels before the ReLU was applied. For the actual and predicted activations in **a-c**, we then constructed a representational similarity matrix (RSM) that contained the same exact conditions as the RSM we measured in the ANN. **d,e)** The hidden layer of the ANN with its corresponding RSM. Note that the RSM of the activity flow-predicted hidden layers was computed in the same way as the ANN – by projecting the input activations of a single condition onto the hidden layer, while fixing all other weights to 0. **f)** The similarity of the actual and activity flow-predicted RSMs with that of the ANN’s RSM. Activations were averaged across subjects first, and then the RSM was constructed using the group-averaged activations for every condition. **g)** Same analysis as in **f**, but computing RSMs for each subject first, and then measuring the similarity of each subject’s RSM with the ANN’s RSM. The similarity of the ANN’s RSM with the activity flow-predicted representations after the ReLU was always greater than the predictions without the ReLU.

**Supplementary Figure 6.**
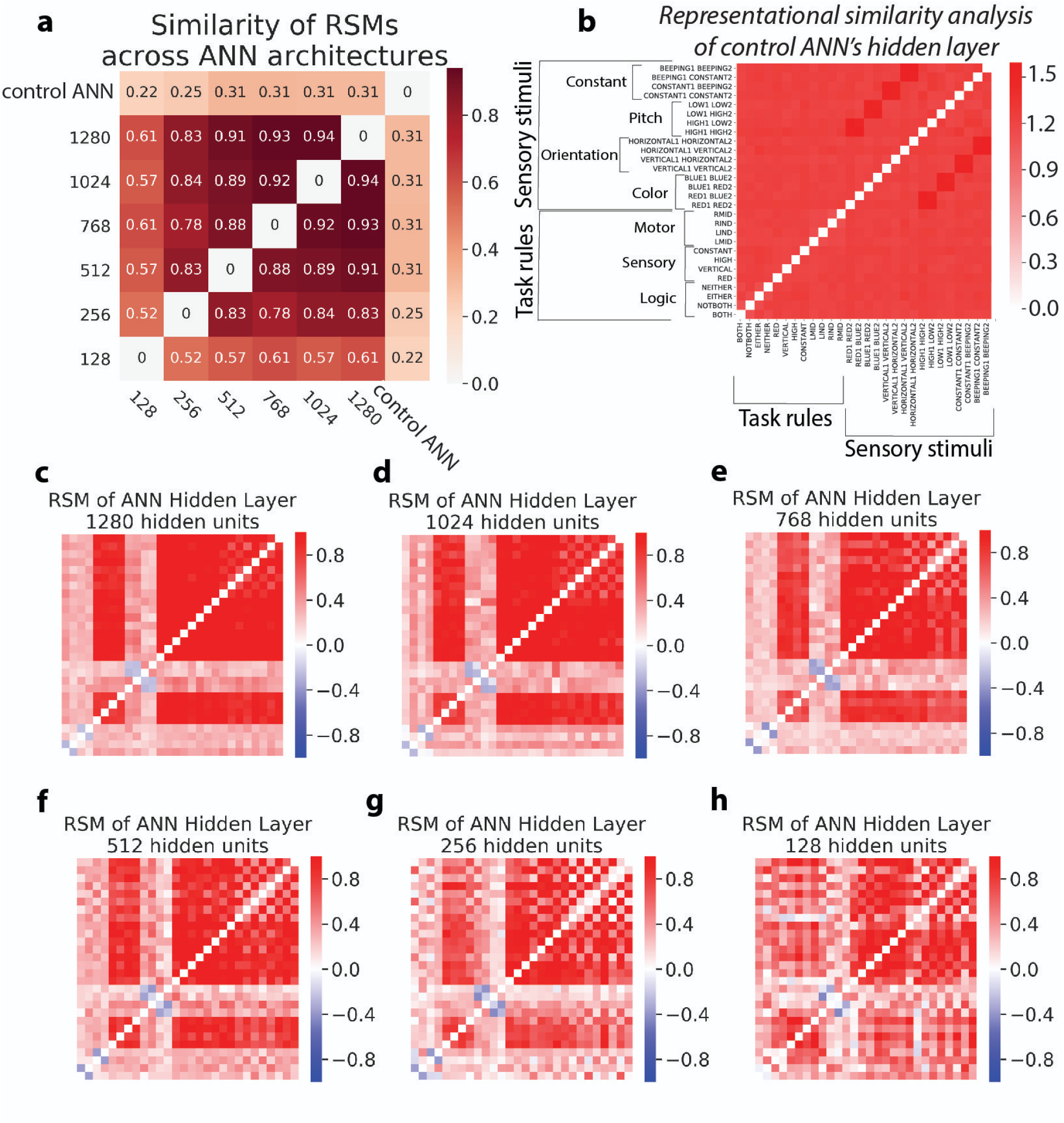
Evaluating the representational similarities of the ANN’s hidden layer across different ANN architectures and controls. **a)** The similarity of the RSMs across different ANNs with different hidden units and the control ANN (i.e., shuffled parameters). We found that ANNs with different numbers of hidden units generally had highly similar representational geometries. However, ANNs with greater hidden units tended to have greater similarity, likely due to improved generalization^68^. **b)** The RSM of the control ANN (shuffled parameters within each layer). **c-h)** The RSMs of ANNs with 1280, 1024, 768, 512, 256, and 128 hidden units. We trained a single instance of each model (trained until accuracy>99.5%) prior to visualizing its RSMs. RSMs with fewer hidden units tended to have more variable (noisier) RSMs across initializationss.

**Supplementary Figure 7.**
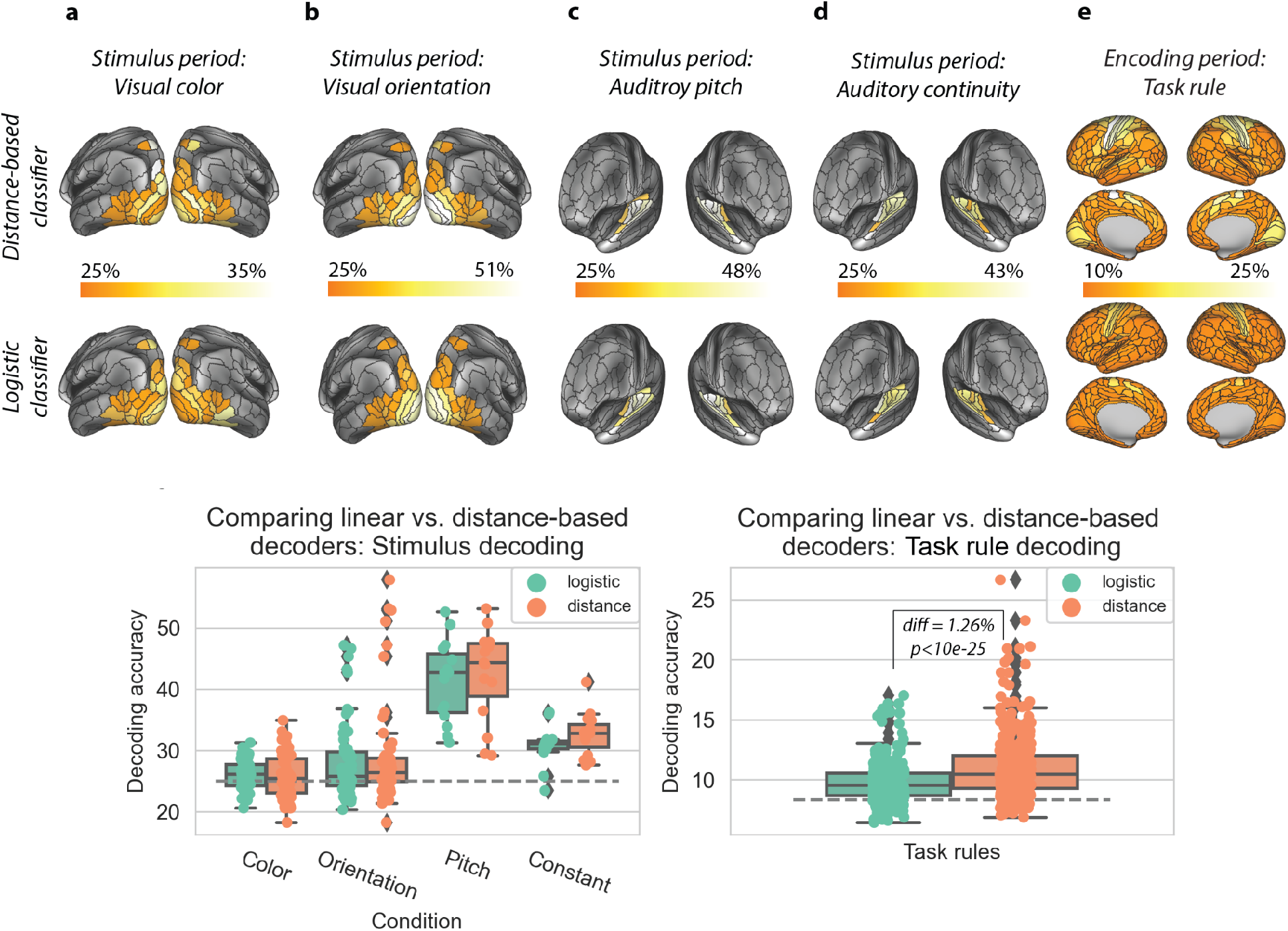
Comparing a linear versus distance-based decoder for identifying regions containing stimulus and task rule features. **a-e)** Surface visualizations for decoding accuracies of each parcel (using vertices within each parcel) for **a)** color (visual), **b)** orientation (visual), **c)** pitch (auditory), **d)** continuity/constant (auditory), and **e)** task rule (12-way decoding). Top surface plots employ the distance-based classifier. Bottom plots employ a logistic classifier. **f)** Comparing overall decoding accuracies for all regions of interests for stimulus features. Plot includes unthresholded classification accuracies. Since stimulus decoding involved a 4-way classification, chance was 25%. Overall, there were no statistically different decoding accuracies for distance-based versus linear logistic decoders. **g)** For task rule classifications, distance-based decoders outperformed logistic classifiers (average accuracy difference +1.26%, p<10e-25 using a Wilcoxon signed-rank test). Overall, distance-based decoders perform similarly or better than logistic decoders for our data set. (Distance-based decoders were also computationally cheaper.)

**Supplementary Figure 8.**
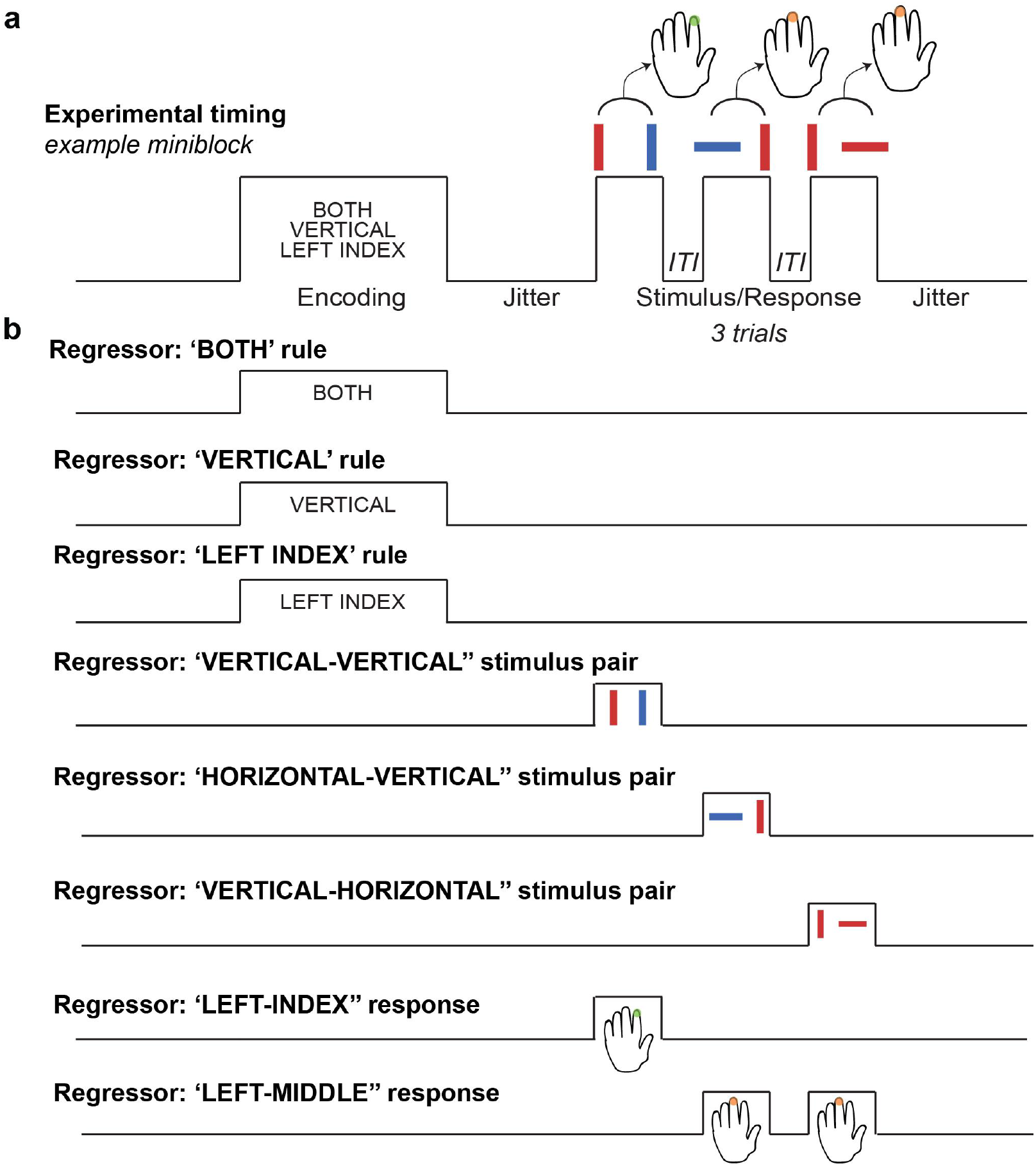
Example of task GLM approach to obtain task activation estimates. **a)** An example miniblock containing one encoding block (task rule set) and three trials. Note that while stimulus presentation and response periods overlap, they are properly counterbalanced across trials. **b)** The regressors for the relevant task conditions in the example miniblock. We obtain regressors (estimated across all 128 miniblocks) for all task rule, sensory stimuli, and motor response conditions. Altogether there are 32 different task conditions (12 task rules, 16 sensory stimuli pairs, and four motor response periods). Note that task rule regressors (logic, sensory, and motor rule examples) appear collinear in this example, but that across all 128 miniblocks task rule conditions are properly counterbalanced to avoid biasing towards any specific rule combination. Regressors shown here are illustrated without convolution with SPM’s canonical HRF.

**Supplementary Figure 9.**
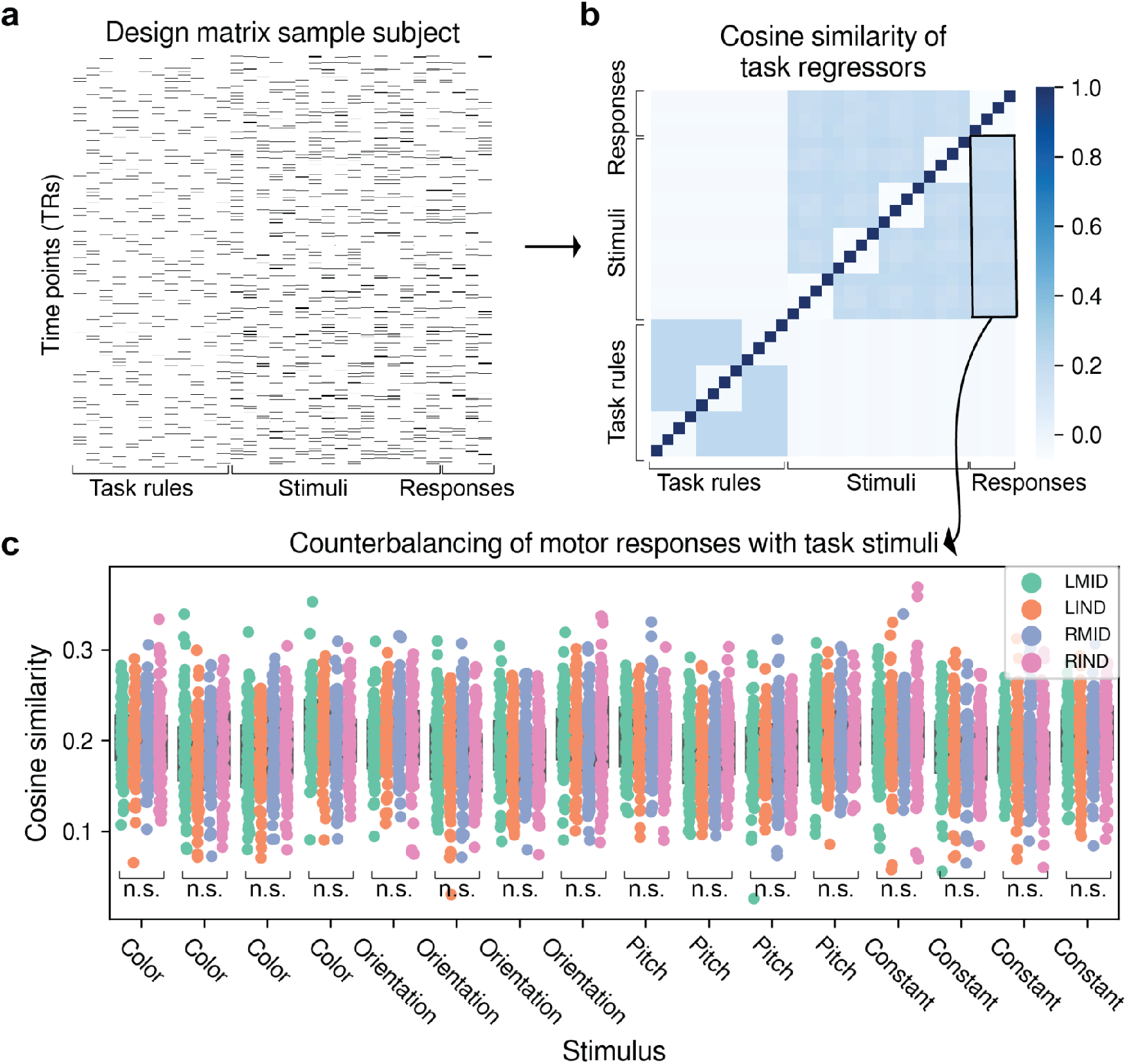
Counterbalancing (and averaging) across conditions ensures statistical independence of activation measures between stimulus and response conditions. **a)** A task GLM design matrix for an example subject. For task GLMs, all conditions (rules, stimuli, responses) were modeled simultaneously using multiple linear regression, increasing the statistical independence of estimates (in addition to counterbalancing). **b)** Group-averaged cosine similarity matrix of task regressors. For each subject, we estimated the cosine similarity of task regressors, then averaged across subjects. Naturally, the diagonal has a cosine similarity of 1. There is no interdependence between task rules and stimulus/motor responses, since the relevant task timing intervals do not overlap. However, for motor and stimulus conditions, there are some weak dependencies (by necessity), since the response intervals overlap with stimulus presentations. However, due to the counterbalancing of stimulus and response types, regressors/conditions will remain independent when modeled together in a task GLM. Therefore, simultaneous modeling of all stimulus and response conditions in a task GLM will ensure unique variance is assigned to each condition. This is because multiple linear regression conditions on all other regressors/conditions. **c)** No main effect of stimulus-response similarity across subjects, demonstrating that averaging across these counterbalanced stimuli eliminates any biases toward particular motor responses. For every stimulus-response pairing, we performed a one-way ANOVA to test the null hypothesis that across response conditions, the distributions had different means (subjects as a random effect). For every stimulus condition, we could not reject the null, indicating no bias towards any given stimulus-response pairing (all p>0.05, FDR-corrected).

**Supplementary Figure 10.**
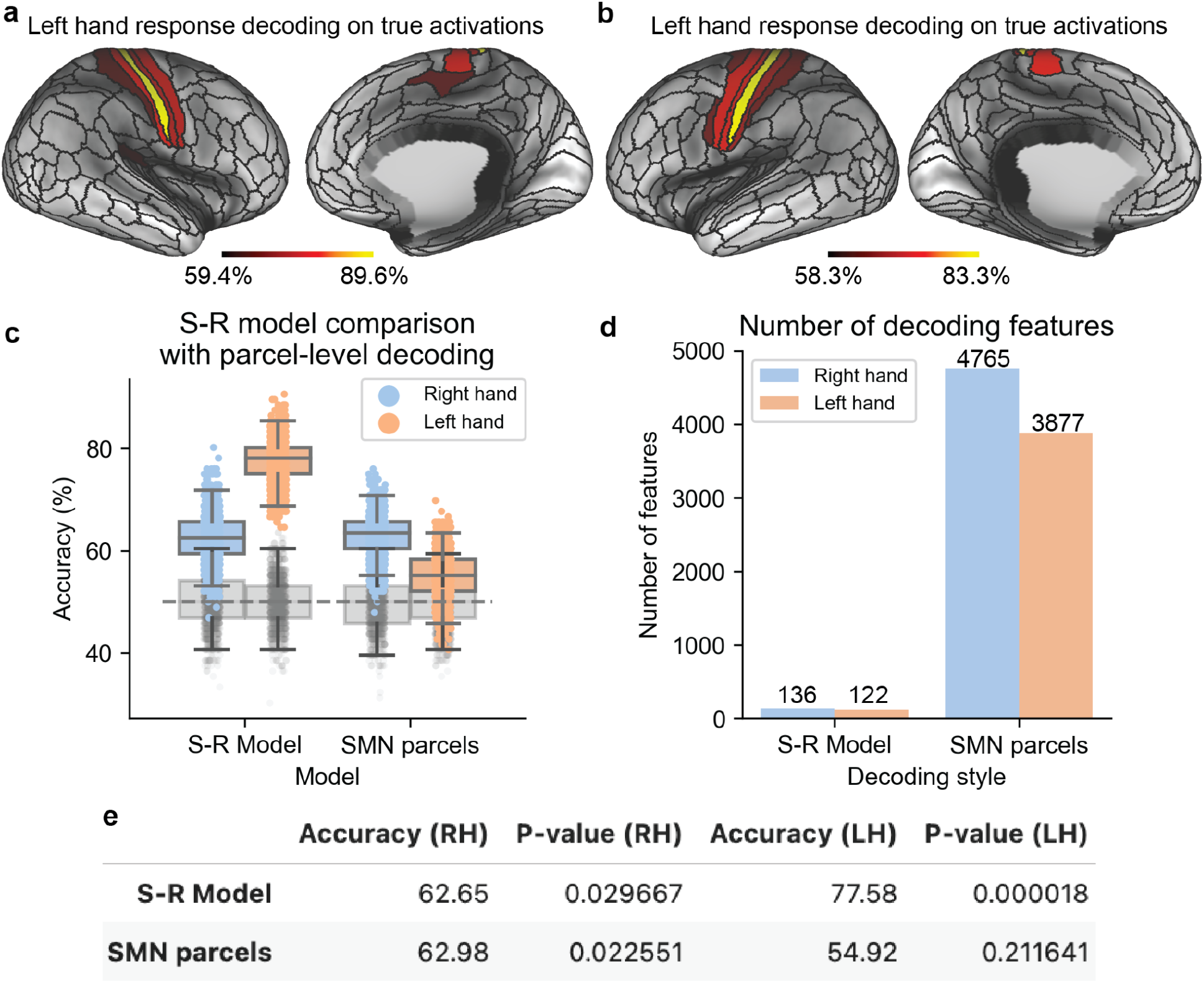
Using classification of response activations to identify motor output parcels reduces spatial specificity of digit representations and reduces stimulus-response decodability for left hand responses. **a)** Left and **b)** right hand response decoding accuracy on the true fMRI response activations. Classifications were limited to the somatomotor network (SMN) on the appropriate hemisphere (e.g., right hemisphere for left hand responses). **c)** S-R model decoding accuracy on the output parcels using the full ENN model (on the left, as presented in the main manuscript; Fig 7b. On the right, the classification accuracy using the SMN parcels identified in **a** and **b**. Right hand classification accuracies remain the same, while left-hand accuracies are degraded. **d)** Number of decoding features that feed the decoder on the output layer. Inclusion of all SMN parcels significantly increases the number of features the decoder is required to distinguish, exacerbating the well-known problem with having substantially more features than observations for classification. **e)** Table of classification accuracies for each model. Cross-validation details are exactly the same as reported in Fig. 7.

**Supplementary Table 1.** Regions containing decodable color (red/blue) stimulus activations.

| Label | GlasserID | Network Affiliation | Hemisphere |
| --- | --- | --- | --- |
| Visual2-54_L-Ctx | L_VVC | Visual2 | L |
| Visual2-05_R-Ctx | R_V4 | Visual2 | R |
| Visual1-03_R-Ctx | R_DVT | Visual1 | R |
| Visual2-22_R-Ctx | R_V4t | Visual2 | R |

**Supplementary Table 2.** Regions containing decodable orientation (vertical/horizontal) stimulus activations.

| Label | GlasserID | Network Affiliation | Hemisphere |
| --- | --- | --- | --- |
| Visual1-04_L-Ctx | L_V1 | Visual1 | L |
| Visual2-28_L-Ctx | L_MST | Visual2 | L |
| Visual2-30_L-Ctx | L_V2 | Visual2 | L |
| Visual2-31_L-Ctx | L_V3 | Visual2 | L |
| Visual2-32_L-Ctx | L_V4 | Visual2 | L |
| Visual2-33_L-Ctx | L_V8 | Visual2 | L |
| Visual2-35_L-Ctx | L_V7 | Visual2 | L |
| Visual2-40_L-Ctx | L_LO2 | Visual2 | L |
| Visual2-41_L-Ctx | L_PIT | Visual2 | L |
| Visual2-42_L-Ctx | L_MT | Visual2 | L |
| Visual2-51_L-Ctx | L_V3CD | Visual2 | L |
| Visual1-01_R-Ctx | R_V1 | Visual1 | R |
| Visual2-03_R-Ctx | R_V2 | Visual2 | R |
| Visual2-04_R-Ctx | R_V3 | Visual2 | R |
| Visual2-05_R-Ctx | R_V4 | Visual2 | R |
| Visual2-06_R-Ctx | R_V8 | Visual2 | R |
| Visual2-09_R-Ctx | R_IPS1 | Visual2 | R |
| Visual2-12_R-Ctx | R_LO1 | Visual2 | R |

**Supplementary Table 3.** Regions containing decodable pitch (high/low) stimulus activations.

| Label | GlasserID | Network Affiliation | Hemisphere |
| --- | --- | --- | --- |
| Auditory-08_L-Ctx | L_A1 | Auditory | L |
| Auditory-09_L-Ctx | L_52 | Auditory | L |
| Auditory-10_L-Ctx | L_RI | Auditory | L |
| Auditory-11_L-Ctx | L_TA2 | Auditory | L |
| Auditory-12_L-Ctx | L_PBelt | Auditory | L |
| Auditory-13_L-Ctx | L_MBelt | Auditory | L |
| Auditory-14_L-Ctx | L_LBelt | Auditory | L |
| Auditory-15_L-Ctx | L_A4 | Auditory | L |
| Auditory-01_R-Ctx | R_A1 | Auditory | R |
| Auditory-02_R-Ctx | R_52 | Auditory | R |
| Auditory-03_R-Ctx | R_TA2 | Auditory | R |
| Auditory-04_R-Ctx | R_PBelt | Auditory | R |
| Auditory-05_R-Ctx | R_MBelt | Auditory | R |
| Auditory-06_R-Ctx | R_LBelt | Auditory | R |
| Auditory-07_R-Ctx | R_A4 | Auditory | R |

**Supplementary Table 4.** Regions containing decodable constant/beeping stimulus activations.

| **Label** | **GlasserID** | **Network Affiliation** | **Hemisphere** |
| --- | --- | --- | --- |
| Auditory-08_L-Ctx | L_A1 | Auditory | L |
| Auditory-09_L-Ctx | L_52 | Auditory | L |
| Auditory-10_L-Ctx | L_RI | Auditory | L |
| Auditory-11_L-Ctx | L_TA2 | Auditory | L |
| Auditory-12_L-Ctx | L_PBelt | Auditory | L |
| Auditory-13_L-Ctx | L_MBelt | Auditory | L |
| Auditory-14_L-Ctx | L_LBelt | Auditory | L |
| Auditory-15_L-Ctx | L_A4 | Auditory | L |
| Auditory-01_R-Ctx | R_A1 | Auditory | R |
| Auditory-02_R-Ctx | R_52 | Auditory | R |
| Auditory-03_R-Ctx | R_TA2 | Auditory | R |
| Auditory-04_R-Ctx | R_PBelt | Auditory | R |
| Auditory-05_R-Ctx | R_MBelt | Auditory | R |
| Auditory-06_R-Ctx | R_LBelt | Auditory | R |
| Auditory-07_R-Ctx | R_A4 | Auditory | R |

**Supplementary Table 5.**
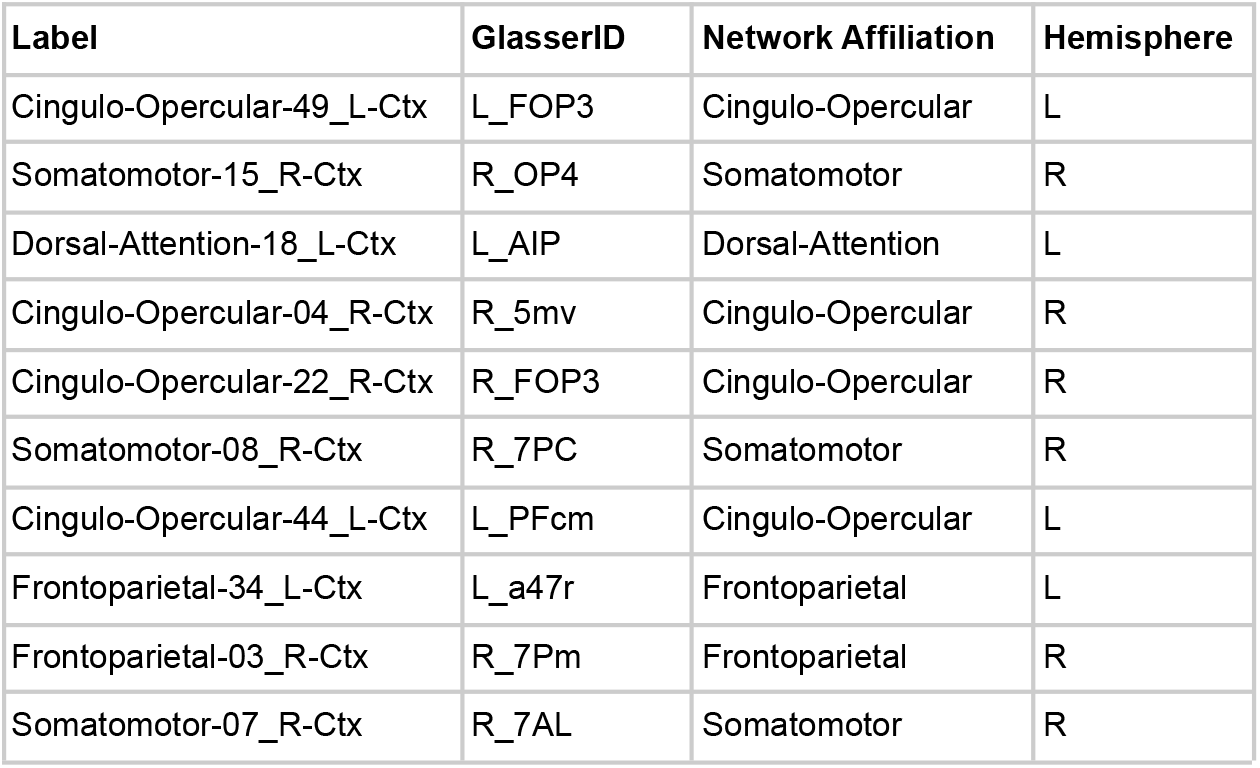
Conjunction hubs.

| Label | GlasserID | Network Affiliation | Hemisphere |
| --- | --- | --- | --- |
| Cingulo-Opercular-49_L-Ctx | L_FOP3 | Cingulo-Opercular | L |
| Somatomotor-15_R-Ctx | R_OP4 | Somatomotor | R |
| Dorsal-Attention-18_L-Ctx | L_AIP | Dorsal-Attention | L |
| Cingulo-Opercular-04_R-Ctx | R_5mv | Cingulo-Opercular | R |
| Cingulo-Opercular-22_R-Ctx | R_FOP3 | Cingulo-Opercular | R |
| Somatomotor-08_R-Ctx | R_7PC | Somatomotor | R |
| Cingulo-Opercular-44_L-Ctx | L_PFcm | Cingulo-Opercular | L |
| Frontoparietal-34_L-Ctx | L_a47r | Frontoparietal | L |
| Frontoparietal-03_R-Ctx | R_7Pm | Frontoparietal | R |
| Somatomotor-07_R-Ctx | R_7AL | Somatomotor | R |

**Supplementary Table 6.**
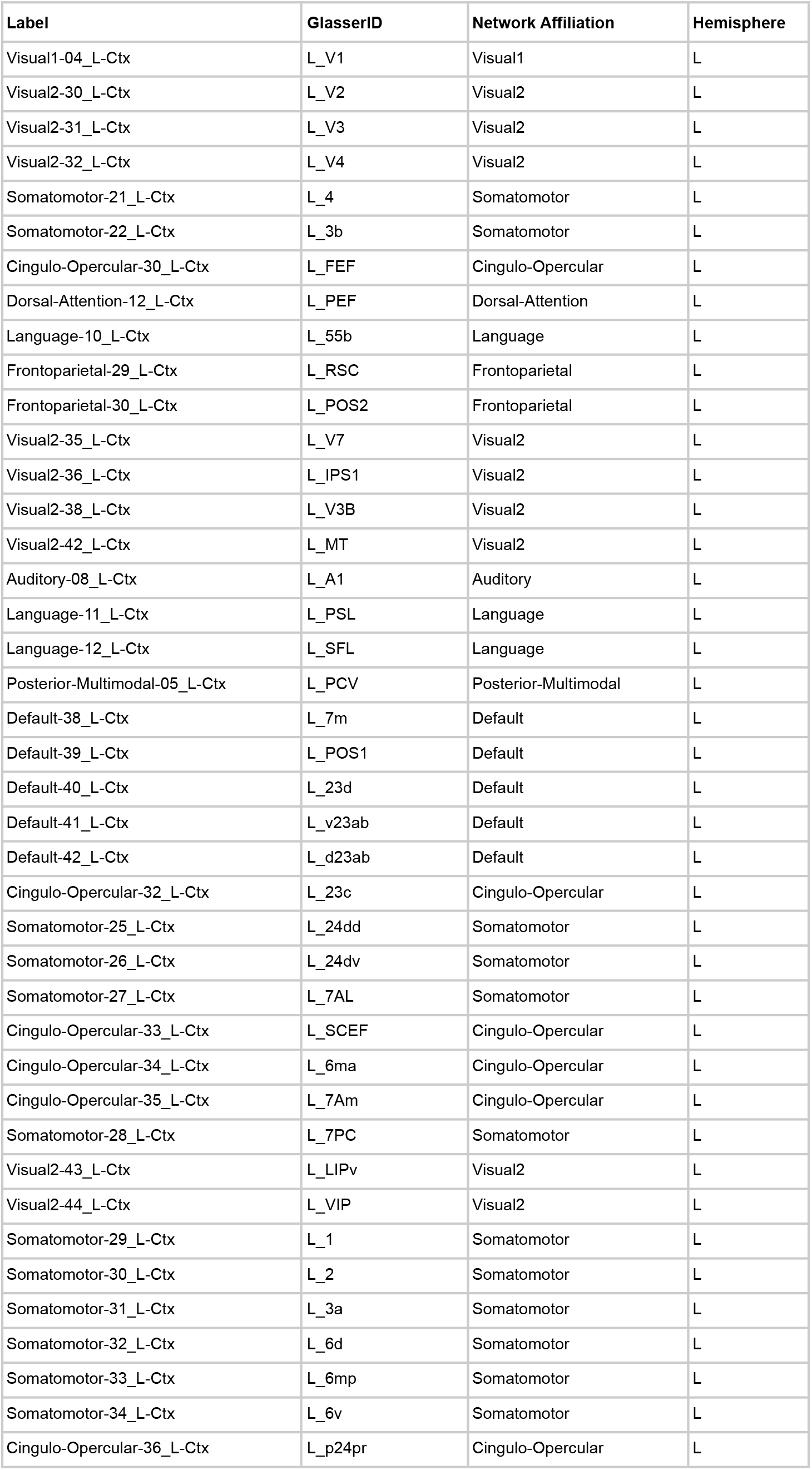
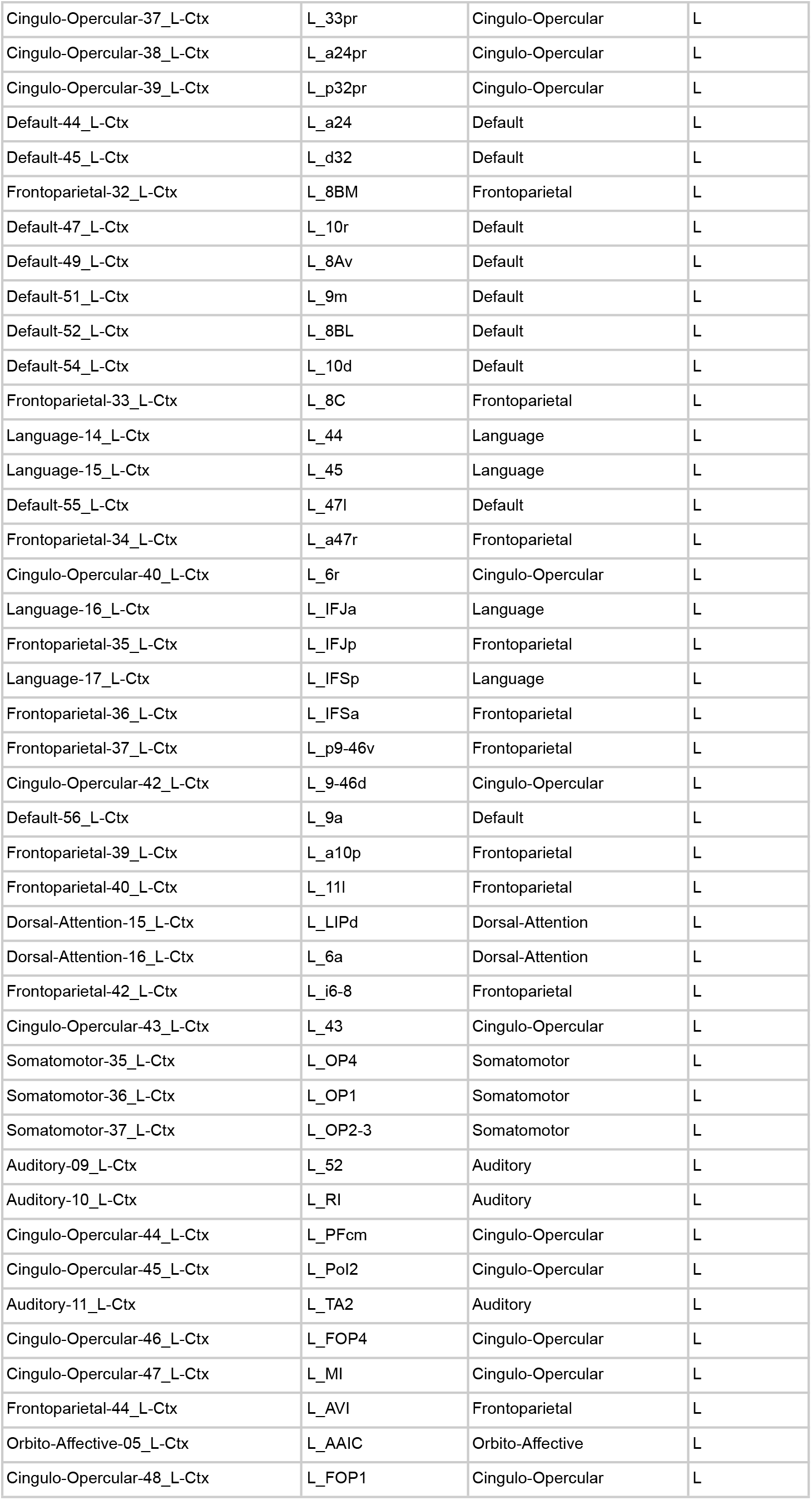
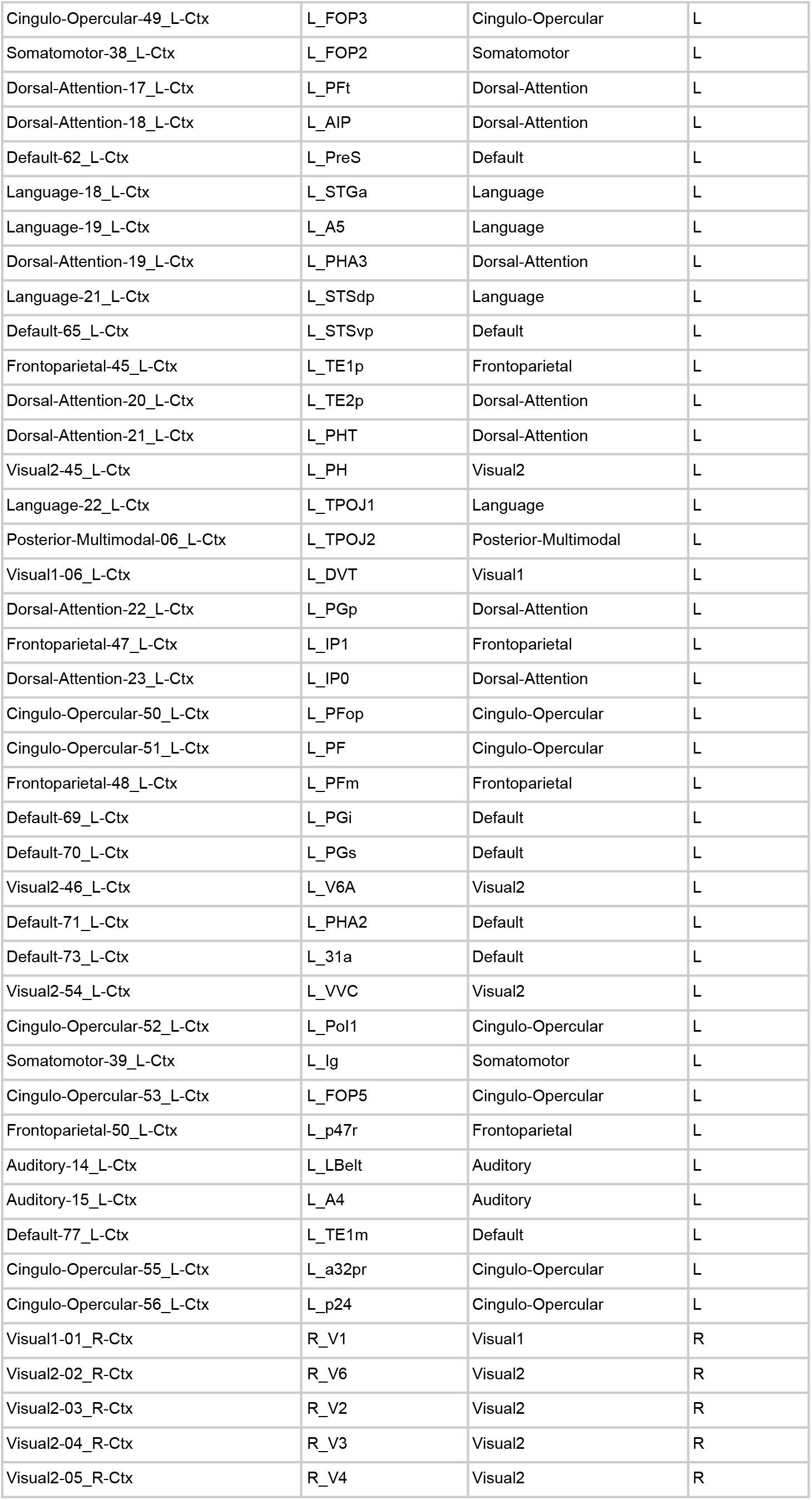
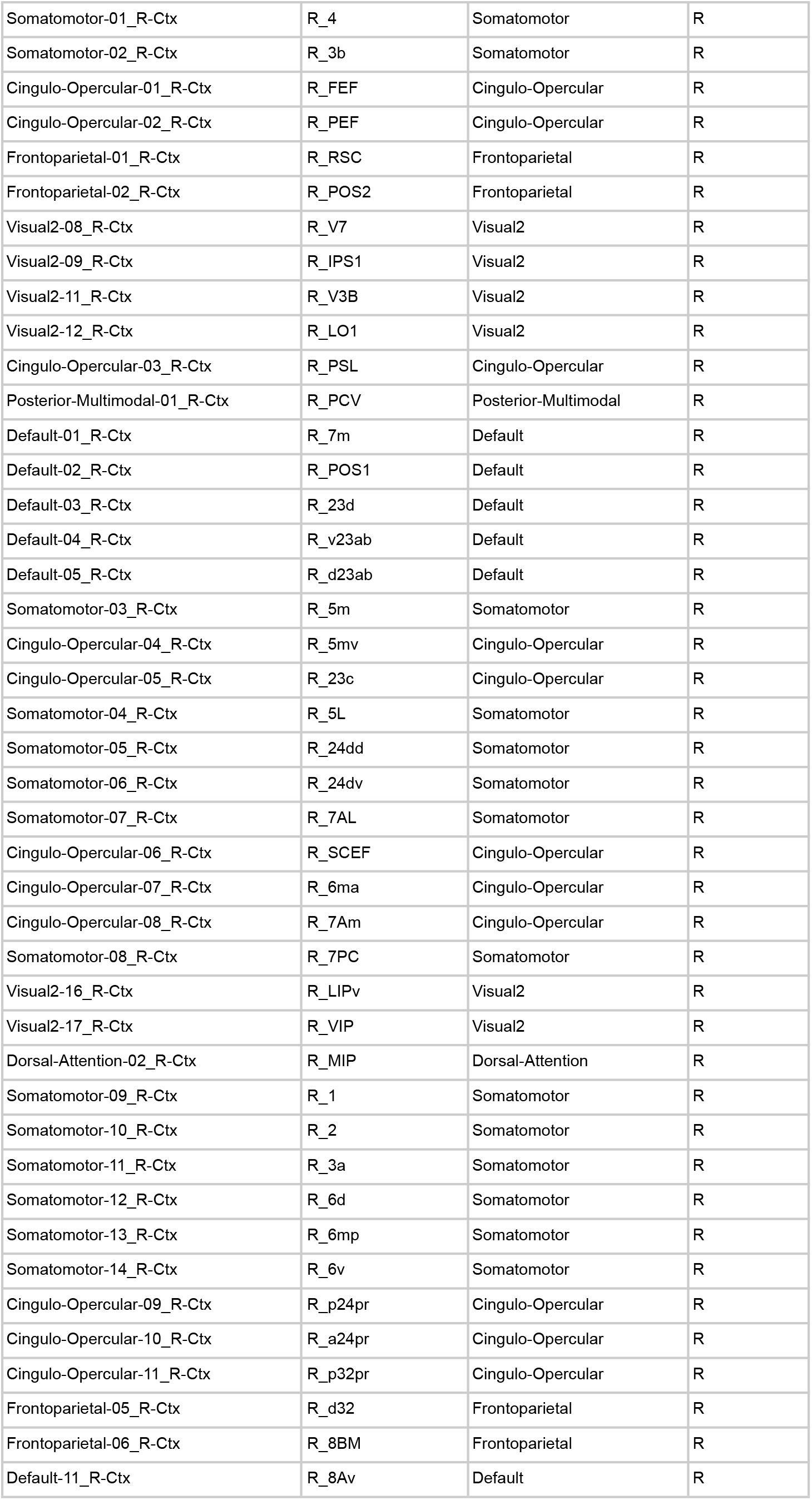
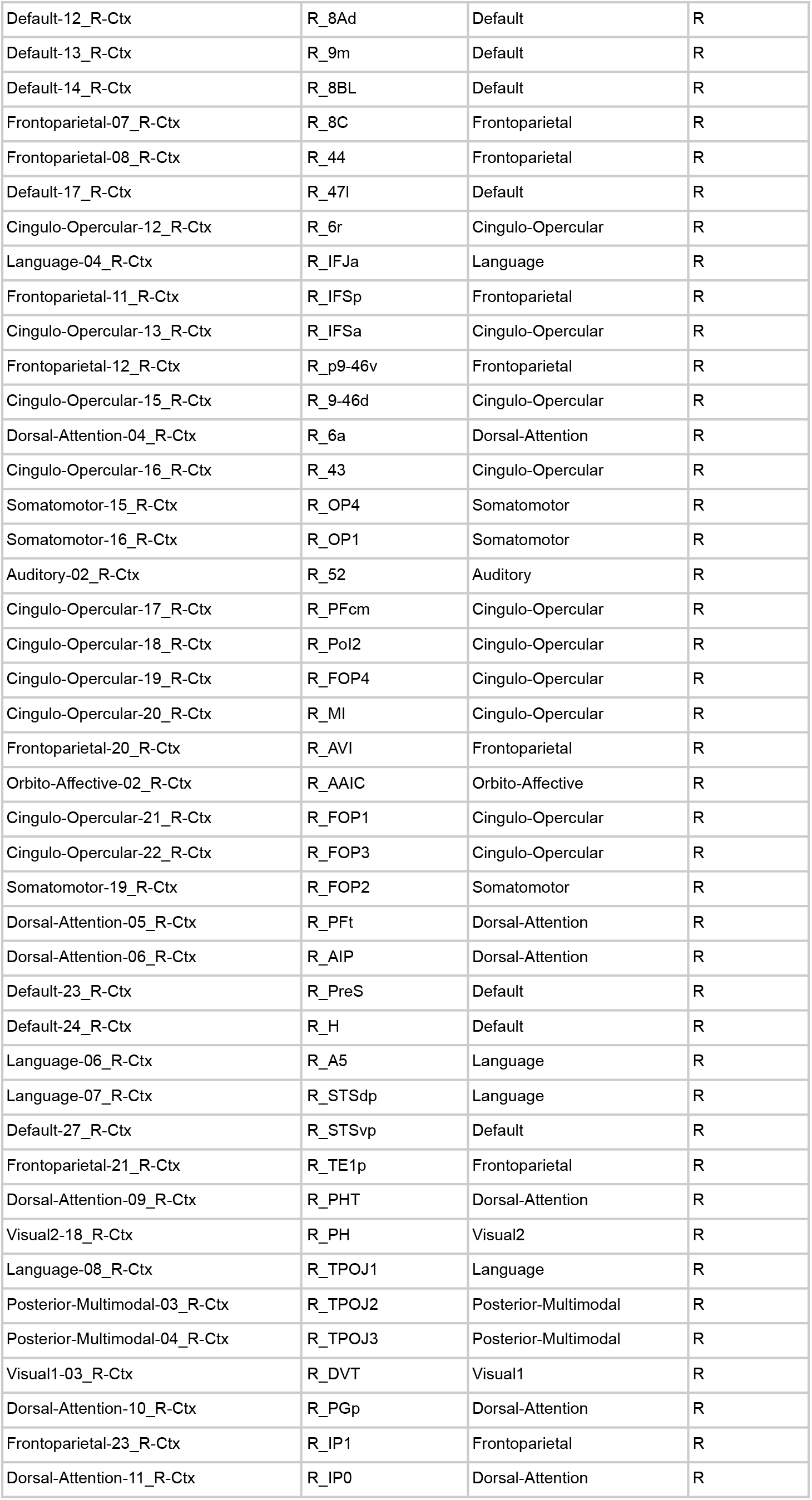
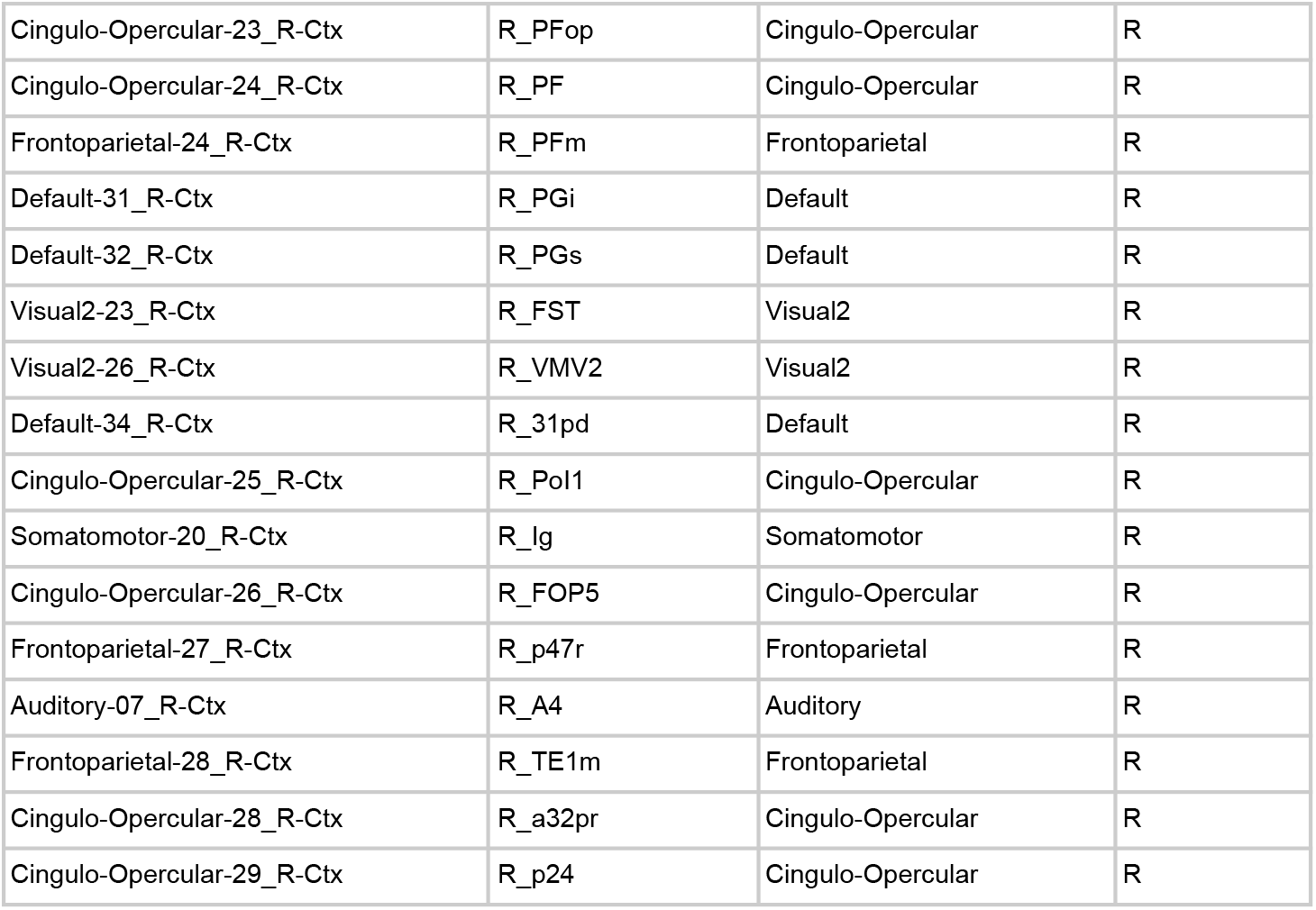
Task rule (input) regions.

